# Integrating multimodal data sets into a mathematical framework to describe and predict therapeutic resistance in cancer

**DOI:** 10.1101/2020.02.11.943738

**Authors:** Kaitlyn Johnson, Grant R. Howard, Daylin Morgan, Eric A. Brenner, Andrea L. Gardner, Russell E. Durrett, William Mo, Aziz Al’Khafaji, Eduardo D. Sontag, Angela M. Jarrett, Thomas E. Yankeelov, Amy Brock

## Abstract

A significant challenge in the field of biomedicine is the development of methods to integrate the multitude of dispersed data sets into comprehensive frameworks to be used to generate optimal clinical decisions. Recent technological advances in single cell analysis allow for high-dimensional molecular characterization of cells and populations, but to date, few mathematical models have attempted to integrate measurements from the single cell scale with other data types. Here, we present a framework that actionizes static outputs from a machine learning model and leverages these as measurements of state variables in a dynamic mechanistic model of treatment response. We apply this framework to breast cancer cells to integrate single cell transcriptomic data with longitudinal population-size data. We demonstrate that the explicit inclusion of the transcriptomic information in the parameter estimation is critical for identification of the model parameters and enables accurate prediction of new treatment regimens. Inclusion of the transcriptomic data improves predictive accuracy in new treatment response dynamics with a concordance correlation coefficient (CCC) of 0.89 compared to a prediction accuracy of CCC = 0.79 without integration of the single cell RNA sequencing (scRNA-seq) data directly into the model calibration. To the best our knowledge, this is the first work that explicitly integrates single cell clonally-resolved transcriptome datasets with longitudinal treatment response data into a mechanistic mathematical model of drug resistance dynamics. We anticipate this approach to be a first step that demonstrates the feasibility of incorporating multimodal data sets into identifiable mathematical models to develop optimized treatment regimens from data.

## Introduction

The development of resistance to chemotherapy is a major cause of treatment failure in cancer. Intratumoral heterogeneity and phenotypic plasticity play a significant role in therapeutic resistance (1,2) and individual cell measurements such as flow and mass cytometry (3) and scRNA-seq (4) have been used to capture and analyze this cell variability (5–8). Although attempts have been made to extract dynamic information from scRNA-seq *via* pseudo-time(9) or RNA velocity approaches (10), these high-throughput “omics” approaches come from cancer cell populations at a single time point. Snapshot information alone has provided immense insight to the field: illuminating novel molecular insight about distinct subpopulations (11), developing detailed hypothesis about population structure (12), and even demonstrating the ability to predict clinical outcomes (1). However, outside of the field of differentiation (13), most “omics”data sets have not been directly integrated with longitudinal population data—which are critical to understanding the dynamics of cancer progression.

Longitudinal treatment-response data in cancer have been used to calibrate mechanistic mathematical models of heterogeneous subpopulations (12,14,15) of cancer cells. These models describe cancer cells dynamically growing and responding to drug with differential growth rates and drug sensitivities. Knowledge of these model parameters have enabled the theoretical optimization of treatment protocols (16–18), and have been applied successfully to prolong tumor control in both mice (12) and patients (14,19). Critical to the success of these modeling endeavors is the ability to identify and validate critical model parameters from available data (20). Identifiable and practical models are necessarily limited in their capacity to explain biological complexity based on the available longitudinal data, which is often limited to total tumor volume or total cell number in time. While we have evidence of complex relationships between distinct subpopulations of cells (11) that give rise to observed behavior, the ability to track these subpopulations longitudinally for use in model calibration and parameter estimation remains a challenge (21).

One way to resolve this challenge would be to work with both types of data (the snapshot “omics” data sets to provide details of distinct subpopulations, and longitudinal population-size data) and use them jointly to inform the calibration of a mechanistic model. In this study, we sought to develop a flexible framework for integrating informatics outputs from high-throughput single-cell resolution data with longitudinal population-size data to demonstrate the feasibility of utilizing multimodal data sources in mathematical oncology. The integration of single cell data into a mathematical modeling framework has been successfully employed in the field of differentiation by quantifying the changing proportion of cells in distinct cell states over time (13). This approach has yet to be applied to cancer, where the effects of exponential growth and death due to drug exposure results in changes in phenotypic composition that are independent of directed transitions between cell states. To better understand these dynamics, we collect longitudinal population-size data in response to treatment with chemotherapy doxorubicin. We combine this with snapshots of lineage-traced scRNA-seq data and build a classifier to estimate phenotypic composition, via the proportion of sensitive and resistant cells, at distinct time points during treatment response. Despite differences in data acquisition, time resolution, and data uncertainty, we demonstrate that these two measurement sources can be used to estimate cell number in time and phenotypic composition in time, which can be compared to their corresponding model outputs. To reflect varying degrees of confidence in the measurement sources, we develop an integrated calibration scheme that relies on Pareto optimality and demonstrate that the phenotypic composition information is essential for the identifiability of model parameters from data. We validate the model results by demonstrating that they can accurately predict the response dynamics to new treatment regimens. We propose this framework as a crucial next step towards combining tumor composition information with longitudinal treatment data to improve prediction and optimization of treatment outcomes.

## Results

### Utilizing a Model of Sensitive and Resistant Subpopulations to Describe and Optimize Drug Response Dynamics

To describe and predict the dynamics of cancer cells in response to treatment, we use a mechanistic model that describes sensitive and resistant cell subpopulations growing, dying, and transitioning from the sensitive, *S*, to resistant, *R*, state as a direct result of treatment (17).

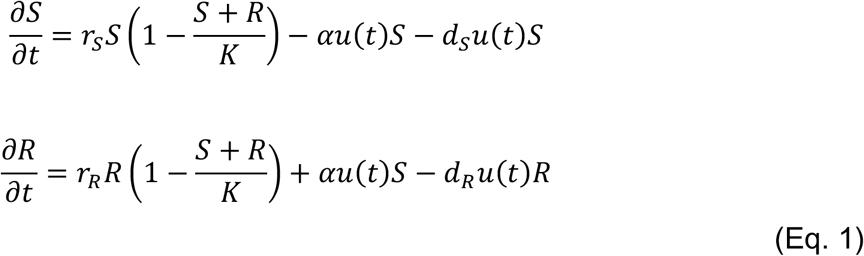

In this model (Fig 1A), sensitive and resistant cells grow *via* a logistic growth hypothesis at their own intrinsic growth rates (*r_S_* and *r_R_*) and a joint carrying capacity (*K*), which will either take the value of *K_N_* for the carrying capacity of the cells in the longitudinal treatment experiment or *K_ϕ_* for the carrying capacity of the cells in the scRNA-seq experiment. Sensitive and resistant cells are killed by the drug at a rate of *d_S_* and *d_R_* respectively, that is proportional to the number of cells in each subpopulation and the effective dose, *u*(*t*), following the log-kill hypothesis. By definition, we set *d_S_* > *d_R_* such that sensitive cells will be more susceptible to death due to treatment than resistant cells. Treatment drives cells from the sensitive subpopulation into the resistant subpopulation at a rate *α*, which is linearly proportional to the number of sensitive cells present and *u*(*t*).

**Fig 1.**
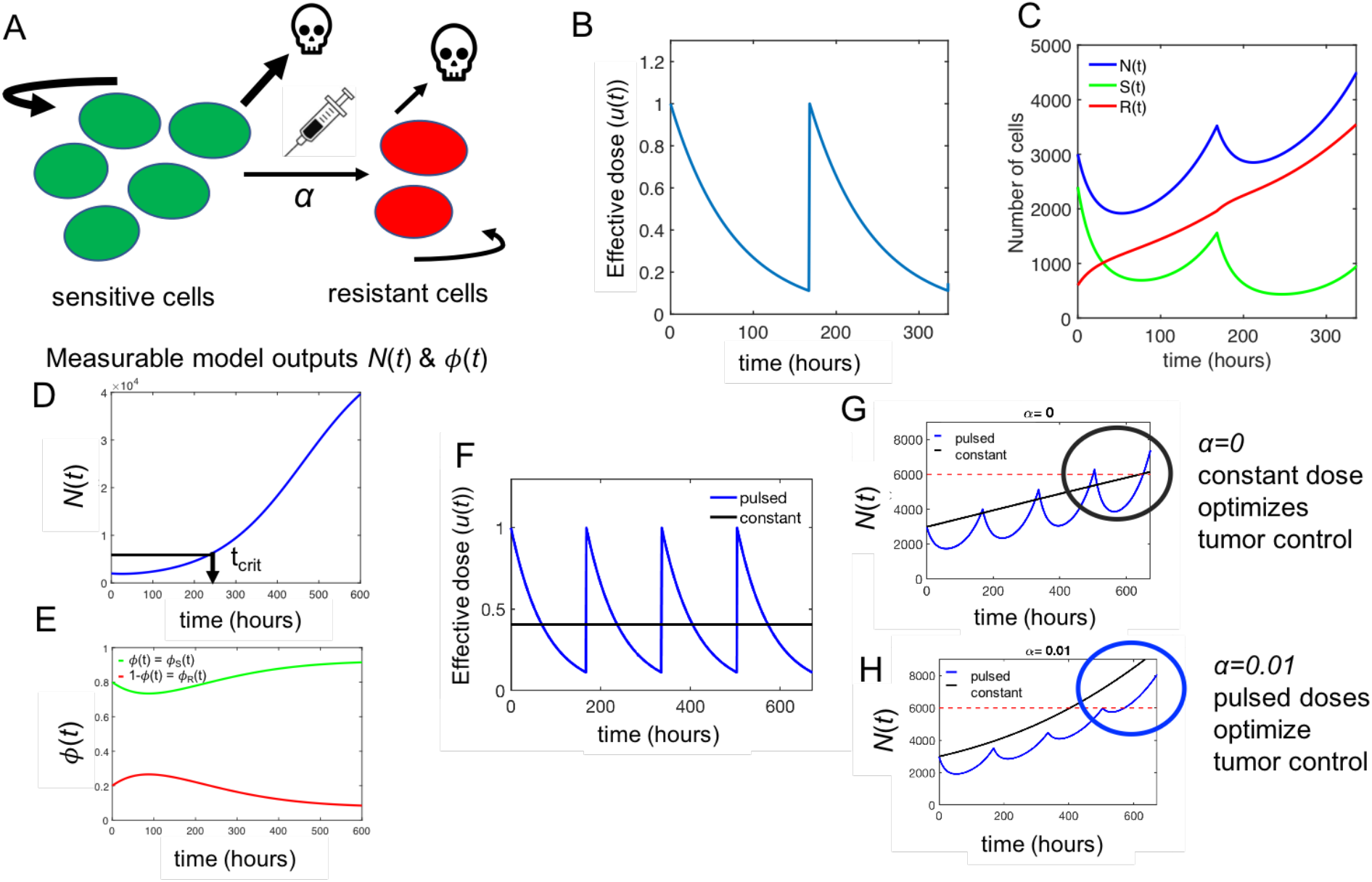
Mathematical Model of Treatment-induced Resistance and its Implications. A. Sketch of the model structure (Eq 1). The model describes sensitive and resistant subpopulations growing exponentially at independent growth rates. In response to treatment, sensitive and resistant cells are killed by the drug. The exposure to drug drives sensitive cells into the resistant phenotype. B. Input effective dose dynamics (*u*(*t*)) for pulse treatment of doxorubicin chemotherapeutic, where exponential decay is assumed (Eq. 2). C. Example of model predicted tumor dynamics under repeated pulse treatments. Sensitive (green) and resistant (red) subpopulation dynamics are predicted by the model. D. Example model predicted total cell number in time in response to a single pulse treatment. The efficacy of a treatment regimen is quantified by the time to reach 2\**N*_0_, which we call *t_crit_* with a longer *t_crit_* indicating a more effective treatment. Experimentally, we can only measure total cell number longitudinally. E. Fraction of cells that are sensitive (green) and resistant (red) in the population over time in response to a single pulse treatment. The phenotypic composition is measured using single cell transcriptomics at discrete time points. F. Pulsed (blue) and constant (black) effective dosing regimens (*u*(*t*)). The constant dose is equal to the average of the pulsed dose over time for ease of comparison (see text for details). G. Example trajectory of total cell number in time for a constant dose (black) and a pulsed dose (blue) for the case where there is no drug-induced resistance (*α* = 0), indicating that optimal tumor control (longer critical time) is reached for the constant dose (black) compared to the pulsed dose (blue). H. Example trajectory of total cell number in time for a constant dose (black) and a pulsed dose (blue) for the case where the drug does induce resistance (*α* > 0), indicating that in this case the optimal tumor control is reached by applying pulse treatments

To investigate the effect of different treatment regimens, we make a simple assumption about the pharmacokinetics of pulsed drug treatments, assuming exponential decay of the effective dose, *u*(*t*), of the drug, as has been shown by others in greater detail (22,23).

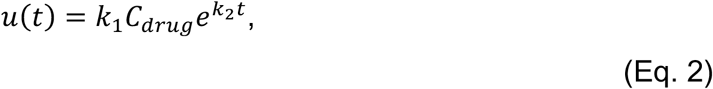

where *C_drug_* is the concentration of doxorubicin in nM, *k_1_* is a scaling factor used to non-dimensionalize the effective dose, and *k_2_* is an estimated rate of decay of the effect of doxorubicin pulse-treatment on breast cancer cells. The effective dose decays over a time scale consistent with experimental measurements of doxorubicin fluorescence dynamics *in vitro* (22,23). An example of the model-predicted treatment response dynamics (Fig 1C) for a pulse treatment given once every week (Fig 1B) demonstrates the response and relapse trajectory in cell number in time (*N*(*t*) in blue), along with the underlying phenotypic dynamics of *S*(*t*) and *R*(*t*). The result of the treatment is that cells in the sensitive population either die or transition to the resistant state, leading to an increase in *R*(*t*) over time even as *S*(*t*) decays and rebounds. For numerical simulations of Eq. 1, we refer to STAR Methods: Model of Drug Resistance Dynamics.

While we can model the dynamics of heterogeneous subpopulations in terms of number of cells in each phenotypic state, most experimental or clinical workflows only allow for measurement of the total cell number (*N*(*t*)) over time (Fig 1D), as single markers of resistance cannot usually be tracked throughout treatment. One unbiased metric to evaluate the response of cell populations to different drug treatments is to measure the time to return to some multiple of the initial cell number (24). We define the critical time (*t_crit_*) as the time it takes for the total cell number to reach double the initial cell number at the onset of treatment (Fig 1D). This metric has been shown to be consistent with “patient benefit” in comparing treatment protocols in pharmacology (24). We employ it here as a single endpoint to evaluate the impact of a treatment on a cell population and to evaluate our model’s predictive capabilities as compared to experimentally measured values of critical time. For a given treatment, while we may not feasibly be able to monitor resistant and sensitive cell number longitudinally, we can estimate the phenotypic composition, which we define here by the sensitive cell fraction, *ϕ_S_*(*t*) (or simply *ϕ*(*t*)), which we will use as a shorthand in the remainder of the manuscript), throughout treatment response from our model (Fig 1E). Model outputs of *N*(*t*) and *ϕ*(*t*) can be used directly to compare to measurements of cell number in time and phenotypic composition in time following a drug treatment. A full description of the parameters in the model system are described in Table 1.

**Table 1.** Description of model parameters to describe resistance dynamics. Descriptions of the parameters either from measured data (Data), fit of the model to the *N*(*t*) (Fit from *N*(*t*)) or *ϕ*(*t*) (Fit from *ϕ*(*t*)), the model assumptions (Fixed), or predicted from the parameter estimation from the fitted model (Predicted). We fit for six free parameters in the calibration scheme, as listed by the first four rows of the table.

| Parameter | Description | Units | Determination |
| --- | --- | --- | --- |
| $r_S, r_R$ | Growth rate of sensitive and resistant cell subpopulations | hour<sup>-1</sup> | Fit from $N(t)$ & $\phi(t)$ data |
| $\alpha$ | Drug-induced rate of transition from sensitive to resistant state | nM<sup>-1</sup> x hour<sup>-1</sup> | Fit from $N(t)$ & $\phi(t)$ data |
| $d_S, d_R$ | Death rate of sensitive and resistant cell populations due to drug, $d_R < d_S$ | nM<sup>-1</sup> x hour<sup>-1</sup> | Fit from $N(t)$ & $\phi(t)$ data |
| $\phi_0$ | Initial proportion of sensitive cells | number of cells | Fit from $N(t)$ & $\phi(t)$ data |
| $K_N$ | Carrying capacity for the longitudinal treatment experiment performed in a 96 well plate to measure $N(t)$ | number of cells | Fit from $N(t)$ untreated control |
| $K_\phi$ | Carrying capacity of the scRNAseq experiment performed in a 10 cm dish to measure $\phi(t)$ | number of cells | Fixed |
| $t_{crit}$ | Time for the number of cells to return to two times the initial cell number ($N_0$) | hours | Data, fit, and predicted |
| $k_1$ | Scaling factor to non-dimensionalize concentration in nM of doxorubicin | nM<sup>-1</sup> | Fixed |
| $k_2$ | Estimated rate of decay of effect of doxorubicin after pulse-treatment | hour<sup>-1</sup> | Fixed |

Previous work has demonstrated the theoretical implications of treatment-induced resistance on identifying optimal treatment regimens (17). Here we also found that for a resistance-preserving therapy (i.e., *α* = 0), a constant dosing regimen optimizes tumor control (black line Fig 1F), leading to a longer critical time than the pulsed treatment (Fig 1G), whereas for a resistance-inducing therapy (i.e., *α* > 0) a pulsed treatment regimen (blue line Fig 1F) optimizes tumor control (Fig 1H). To compare the effects of a constant versus pulsed dose, we simulated the effect of a constant dose (black line Fig 1F) equal to the mean value over the time interval simulated of the pulsed dose (blue line Fig 1F) in an attempt to reflect realistic toxicity constraints that would be present in a clinical setting when developing treatment regimens. This analysis, as well as further work to utilize this modeling framework to develop optimal treatment protocols (16) indicates that identifying these model parameters is essential to implementing more sophisticated treatment strategies in a practical clinical setting. While (16) show that the critical model parameters are theoretically structurally identifiable from population size data alone, we seek to demonstrate how this model can be practically identified from *in vitro* data using both longitudinal population size data (*N*(*t*)) and snapshot outputs of the phenotypic composition (*ϕ*(*t*)) at a few time points, enabled by recent advances in lineage tracing (11,25) and scRNA-seq technologies. We present this project workflow in the experimental setting as proof-of-concept of the ability to properly identify key model parameters from multimodal data sets, with the hopes that the approach of integrating snapshot with longitudinal data sets will eventually be brought to the clinic to develop optimized treatment regimens for existing therapeutic agents.

To demonstrate the feasibility of integrating multimodal data sources into a cohesive modeling framework, we employ an experimental *in vitro* model system of MDA-MB-231 triple negative breast cancer cells exposed to the chemotherapeutic doxorubicin. The combined experimental-computational workflow (Fig 2) starts by tagging individual cells with unique barcodes that are integrated into the genome and expressed as sgRNA’s; this COLBERT cell barcoding platform has been described previously (25). The barcode-labeled cell population is expanded to generate the naïve population for these studies (305 unique barcodes represents 305 clonal subpopulations). Cells are then treated with doxorubicin (LD95, 550 nM) for 48 hours and allowed to recover; scRNA-seq is performed prior to treatment and from two parallel replicates after the population had regrown following the pulse treatment.

**Fig 2.**
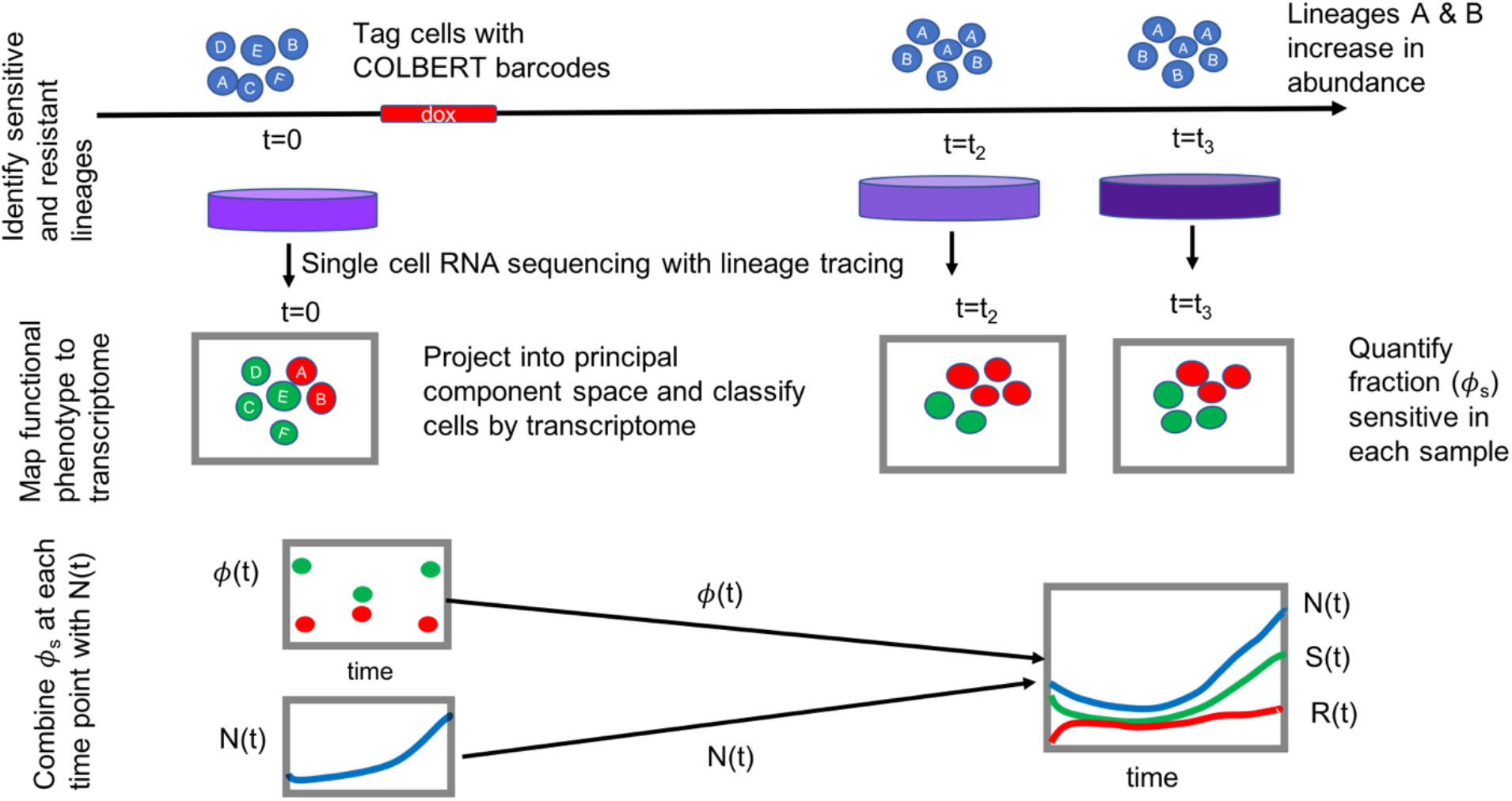
Schematic of the workflow for identifying model parameters from data. At *t* = 0 prior to treatment, individual cells are tagged with a unique, heritable, expressed COLBERT barcode. Cells are treated with a pulse treatment of doxorubicin and allowed to recover from treatment, at which time the barcode abundance is quantified. Lineages whose barcode abundance increased from pre-to post-treatment are assumed to have been in a phenotypic state at *t* = 0 that conferred them more resistant to drug than cells whose barcodes significantly decreased in abundance after treatment. Samples of the population were taken before and from parallel replicates sampled at two different time points after treatment for scRNA-seq. The transcriptomes in the pre-treatment samples of the cells tagged with resistant lineages are assigned resistant and the cells tagged with sensitive lineages are assigned sensitive. Using the gene-cell matrix and labeled class identities of sensitive or resistant from the pre-treatment time point only, a classifier is built using Principle Component Analysis (PCA) to distinguish between sensitive and resistant cells. The classifier is applied to the remainder of transcriptomes of the cells, resulting in a prediction for each cell as either sensitive or resistant. These machine learning outputs are made actionable as state variables by using them to quantify the proportion of sensitive cells (*ϕ*(*t*)) at the three time points. This is combined with separate experiments of longitudinal treatment response dynamics (*N*(*t*)) of the bulk population of the same cell type, and both serve as measured data to be compared to model predicted outputs for parameter estimation.

The transcribed barcode sequence is measured with other transcripts in scRNA-seq. Clones which significantly increase in abundance after treatment are labeled as resistant and those which decrease are labeled as sensitive. We then map the resistant and sensitive functional phenotypes to the transcriptomes of the individual cells they correspond to at the pre-treatment time point. A machine learning classifier is built based on the labeled cell identities and their transcriptomes, and we can apply this classifier to each of the “unknown” cell identities (phenotypes) from the remaining samples. Estimating the binary phenotype of each individual cell from a sample taken throughout treatment response, we quantify the phenotypic composition (*ϕ*(*t*)) at each time point that scRNA-seq was performed. This phenotypic composition measurement can then be combined with longitudinal population size data from drug treatments at different concentrations, compared to corresponding model outputs, and serve to calibrate the mathematical model of drug-induced resistance (Supp Fig. S1).

### Lineage-Traced scRNA-seq Enables Identification of Sensitive and Resistant Phenotypes

To investigate the dynamic changes in phenotypic composition in response to treatment, we sought to characterize gene expression over time at the single-cell level. ScRNA-seq was performed on a barcode-tagged cell population at three time points: immediately before treatment and at parallel replicate samples taken at 7 and 10 weeks after doxorubicin treatment (see STAR Methods: Integration, expression, and capture of COLBERT barcodes). By quantifying the proportion of cells with each lineage identity before and aggregated after treatment, we could identify a functional phenotype associated with the pre-treatment transcriptomes. We quantified changes in lineage abundance (percent of the post-treatment population minus percent of pre-treatment population) of each lineage present in the pre-treatment sample to obtain a distribution of changes in abundance after treatment (Fig 3A). Cells whose lineage abundance increased by any amount after treatment were labeled resistant, and cells whose lineage abundance decreased by more than 5% were labeled sensitive (Fig 3A). All other cells remained unlabeled. This resulted in a training set of 815 labeled cells and their expression levels of 20,645 genes.

**Fig. 3.**
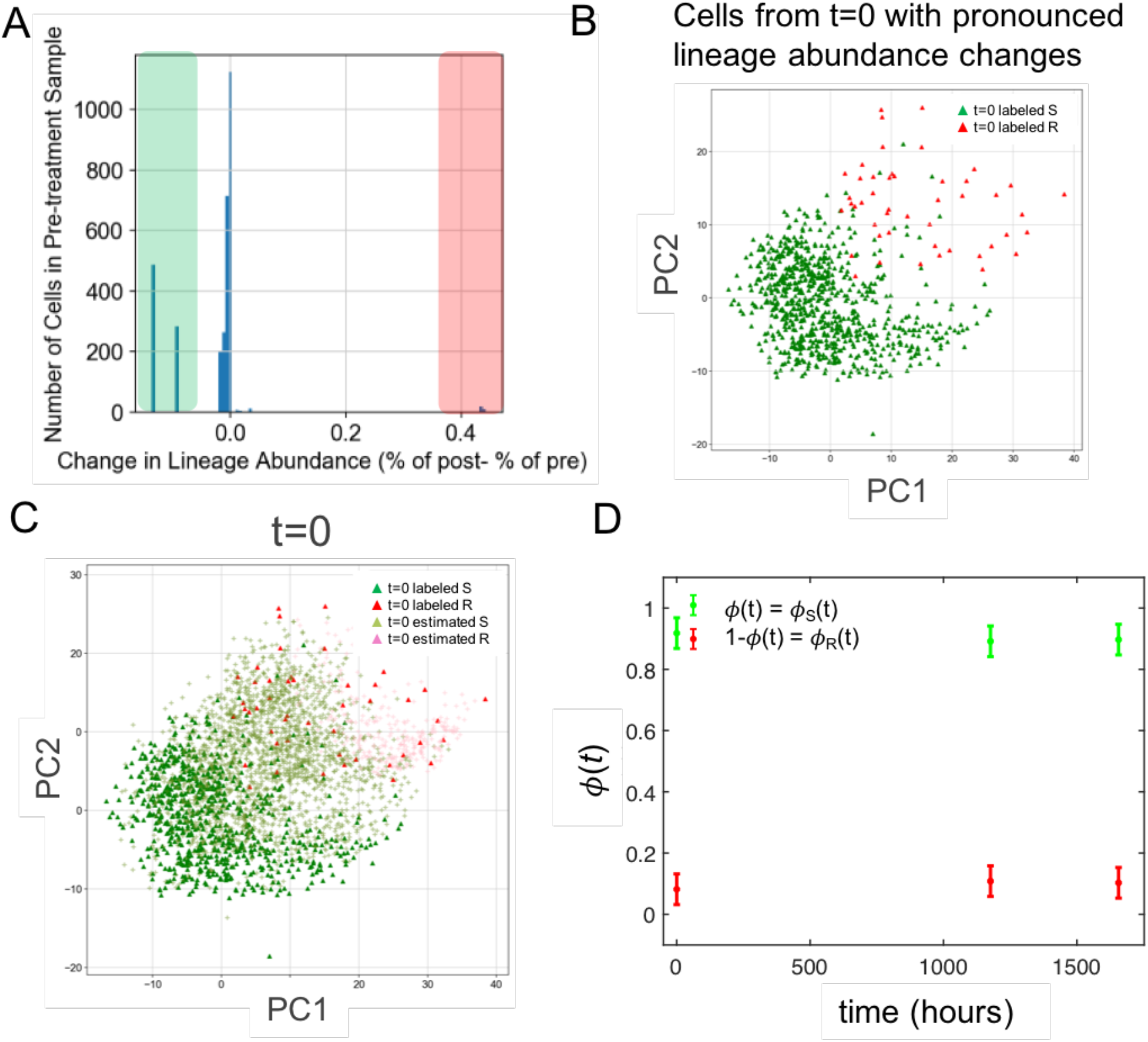
Functional Read-out of Changes in Lineage Abundance Allows Mapping of Phenotypes to Transcriptome. A. Distribution of changes in lineage abundance from pre-to post-treatment indicates separation of lineages whose cells survive and proliferate and those that are more likely to have been killed by the drug treatment. B. Cells from the extremes of high and low lineage abundance changes (highlighted in green and red in A), projected into principal component space display separation along components (cells are projected into PC1 and PC2 space for visualization, full PC-space is made up of 500 principal components, from the initial 20,645 genes detected). C. Example of remaining cells from *t* = 0 projected onto labeled cells in PC space and estimated as sensitive (olive) or resistant (pink). This was performed for the remaining two time points as well (*t* = 7 weeks and *t* = 10 weeks) D. Proportion of cells classified as sensitive or resistant at each time point is quantified from each samples projection and classification as is displayed in C.

The cells from the identified lineages in the pre-treatment time point were labeled as sensitive or resistant as described above, and the labeled gene-cell matrix was used to build a classifier (based on principal component analysis) capable of predicting whether a cell is more likely to be in a resistant or sensitive state based on its gene expression information alone. See STAR Methods: Machine Learning of Cell Phenotypes for full description of building of the classifier. The optimal hyperparameters of the number of nearest neighbors and the number of principal components for class separation were determined based on 5-fold cross validation and coordinate optimization and were found to be 500 principal components and 73 nearest neighbors (Supp Fig. S2). A full description of the methods for hyperparameter optimization are outlined in the STAR Methods: Hyperparameter Optimization. Labeled cells are projected into the principle component space, as is displayed visually using projections into only PC1 and PC2 in Fig 3B, for sensitive cells (green) and resistant cells (red). To identify the phenotype of new cells that the classifier is not trained on, we project each cell into the principle component space of the labeled cells. A k-nearest neighbor graph is constructed to identify the class of the 73 nearest neighbors in the space, and these are averaged to find the probability of the new cell being in the sensitive or resistant state, where cells above a probability threshold are estimated as sensitive (olive) and below the threshold are estimated as resistant (pink) (Fig 3C). This is done for the remaining pre-treatment samples as well as for the cells from the 7 week and 10-week time points (Supp. Fig S3) based on the single cell gene expression vectors (transcriptomes) of each individual cell. We use the machine learning output to predict each cell as either sensitive or resistant and make these predictions actionable by leveraging them as measurements of state variables, the proportion of sensitive cells over time *ϕ*(*t*) (Fig 3D). This quantity will be used as one measurement source for model calibration (Eq. 1) (Supp Fig. S1A & B)

### Ability to Classify Cells as Treatment Sensitive and Resistant Enables Mechanistic Insight into Hallmarks of the Resistant Phenotype

Having identified cells as either resistant or sensitive based on a functional read-out of post-treatment abundance, we can use the class estimates to better understand the transcriptional differences between resistant and sensitive cells. Because the cells largely overlap in principal component space, we use Uniform Manifold Approximation and Projection (UMAPs) as an alternate dimensionality reduction technique for visualization only of the scRNAseq data from the three time points (See STAR Methods: Single Cell Normalization). UMAP projections allow for separation of the three time points (Fig 4A), and within this projection we can highlight which cells were estimated as sensitive (green) and which cells were estimated as resistant (red) (Fig 4E). We can see that the resistant and sensitive cells separate along the first two principal components from Fig 4B, where resistant cells accumulate in the upper right quadrant (high in PC1 and PC2) and sensitive cells aggregate in the lower left quadrant (low in PC1 and PC2). Although these principal components only make up a small proportion of the observed variance in gene expression (Supp Fig. S2D), we see a significant drop off in observed variance after the first few components, indicating that the weights of the genes in these first two components can likely provide us with some mechanistic insight as to which genes are most highly weighted in determining drug-resistance classification. We select a subset of the gene loadings and plot their direction in the first two principal components (Fig 4C), along with a heatmap of the average expression levels for the sensitive and resistant cells for each time point for the top 50 weighted genes in PC1 (Fig 4D). The heatmap reveals that the patterns of regulation between genes in the sensitive and resistant cell classes are conserved across the time points. Examples of the differential expression of NEAT1, UBE2S, and TOP2A are shown in Fig 4 F, G, and H respectively. Comparing these gene expression maps to the UMAP of resistant and sensitive cell classes (Fig 4E) we can see that increased expression in NEAT1 is associated with resistance, while increased expression in UBE2S and TOP2A are associated with the sensitive state. Mechanistically, this corroborates previous findings, as NEAT1 has been shown to be associated with resistance in triple negative breast cancer (26). Although we do not perform detailed molecular analysis in this work, the framework presented here to distinguish sensitive and resistant cells over time can be used to perform a more detailed mechanistic investigation of molecular drivers of resistance, and that is an area of future work. For now, we present the results to demonstrate the interpretability of the classifier and its ability to be validated by examining the gene expression levels of known markers of resistance in the components of the classifier.

**Fig 4.**
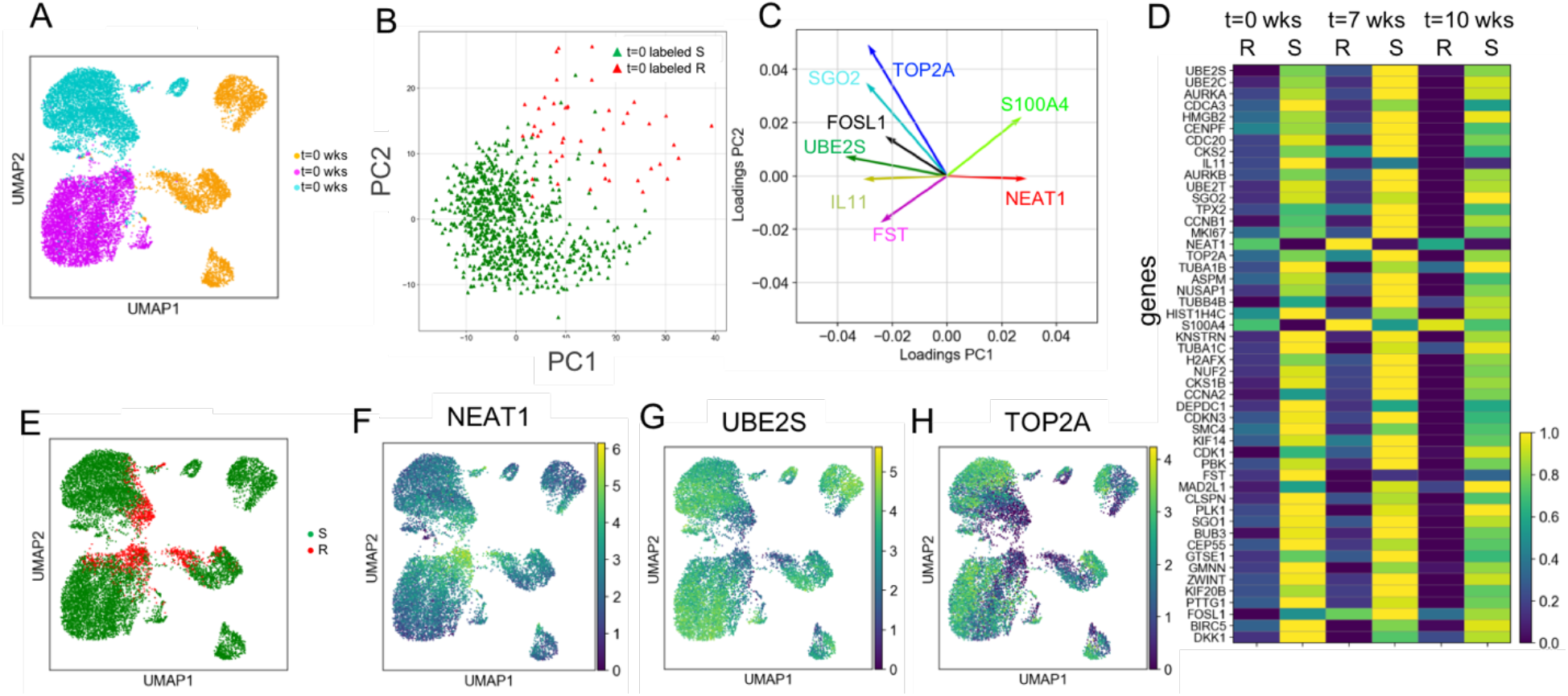
Principal Component Analysis and Differential Gene Expression Analysis Provide Molecular Insight into Drug Resistance Interactions. A. UMAP projection of single cell transcriptomes colored by time point. B. Labeled sensitive and resistant cells projected into the space visualized by the first two principal components, PC1 and PC2, indicating that resistant cells cluster in the upper right quadrant (high in PC1 and PC2), and sensitive cells tend to cluster in the bottom left quadrant (low in PC1 and PC2). C. Gene loadings for selected genes plotted in the space of the first two principal components illuminates key genes that may be associated with resistance to doxorubicin. D. Heat map of the top 50 genes in PC1 comparing the average expression across the sensitive and resistant cell groups in the three time points, showing a characteristic pattern between sensitive and resistant cells across the three time points. The color bar is scaled within each gene (row). E. Single cells colored by sensitive (S) and resistant (R) cell classifier labels visualized via UMAP projections indicates drug sensitivity phenotypes cluster together, but not exclusively by the apparent UMAP clustering. F. UMAP projections of cells colored by expression level of NEAT1 indicates high expression of NEAT1 is associated with resistance. G. UMAP projections of cells colored by expression level of UBE2S indicates that high expression of NEAT1 is associated with sensitivity. H. UMAP projections of cells colored by expression level of TOP2A indicates that high expression of TOP2A is associated with sensitivity.

### Experimental Measurements of Population Size Dynamics in Response to Varying Pulse Treatments of Doxorubicin

In any attempt to model changes in subpopulation frequencies in cancer, bulk population dynamics reflecting differential growth rates and drug sensitivities need to be taken into account. In pivotal work by (13) in the field of differentiation, the proportion of cells in three well-defined differentiation states was used to calibrate mathematical models to describe the mechanism of directed transitions in the differentiation process. Over this time scale, it could reasonably be assumed that no significant differential growth rates accounted for changes in composition. However, in cancer and specifically in this study, we monitor populations over much longer time scales and it is necessary to also consider the contribution of differential growth and death rates among subpopulations. This requires measurements of both bulk population dynamics and subpopulation frequencies over time. In the experimental *in vitro* setting, quantifying bulk population size dynamics is quite feasible for a range of treatment conditions, allowing us to observe the differences in response due to various drug concentrations. Cells were treated with a 24-hour pulse of one of 10 doxorubicin concentrations (ranging from 0-1000 nM, *n*= 6 replicate cell populations for each dose) (Fig 5A, 5B) and the cell number monitored throughout regrowth by time-lapse microscopy. The mean and 95% confidence intervals of cell number in time are shown in Fig 5C. For each dose, we measured the critical time, *t_crit_*, on this measurement of treatment efficacy (Fig 5D). As expected, the higher the dose, the longer the population remained below the critical cell number. The cell number in time data, *N*(*t*), will be used as another measurement source for model calibration (Eq.1) (Supp Fig S1C&D).

**Fig 5.**
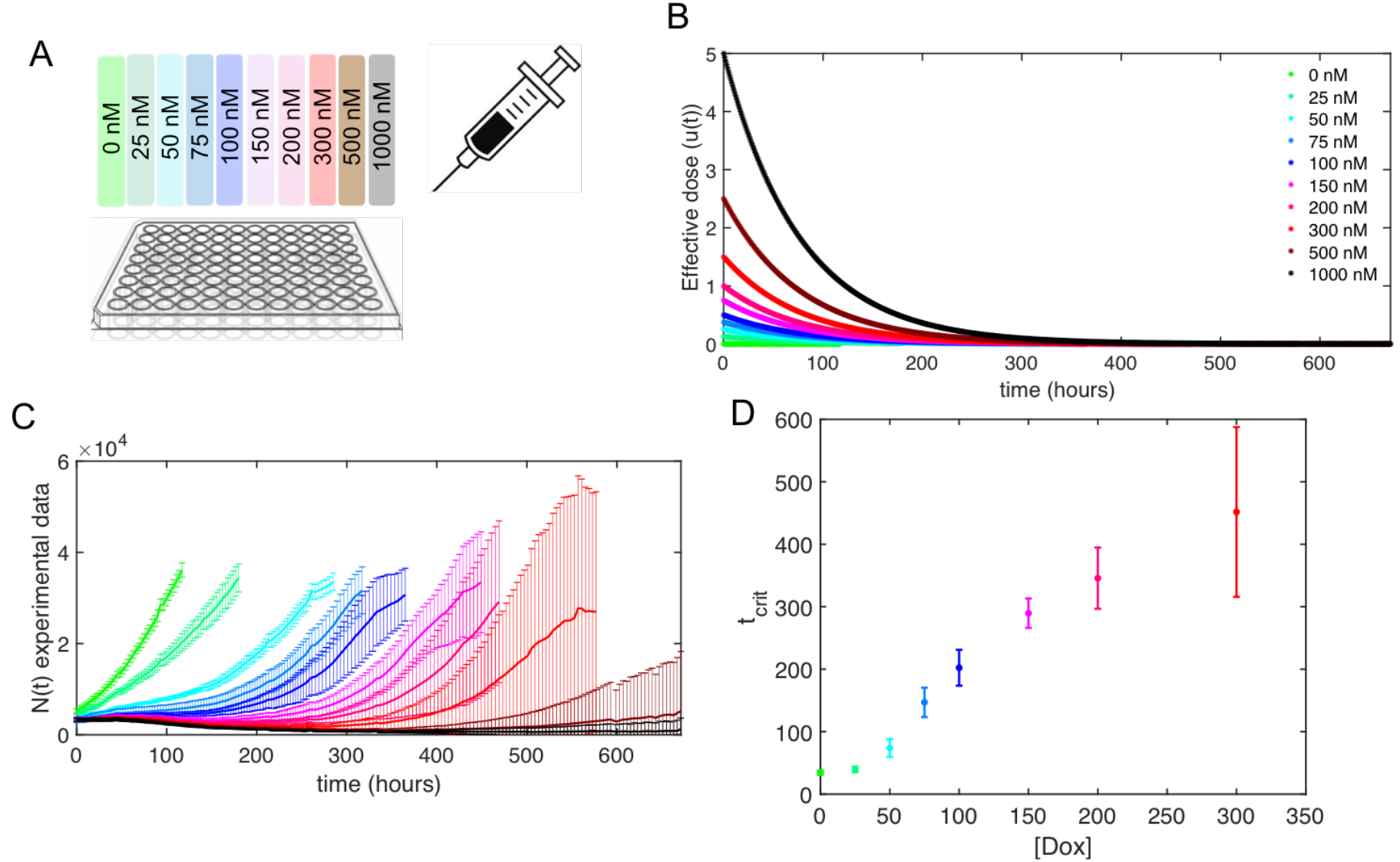
Longitudinal Treatment Response Dynamics for Pulse Treatments at Ten Different Drug Concentrations. A. Schematic of experimental set-up using time-resolved fluorescence microscopy to measure the number of MDA-MB-231 GFP labeled breast cancer cells in response to doxorubicin concentrations ranging from 0-1000 nM treated for 24 hours and then allowed to recover in growth media. B. Estimated effective dose dynamics (*u*(*t*)) of the various pulse-treatments of doxorubicin. C. Number of cells in time, colored by drug concentration as in B, from six replicate wells. Error bars represent 95% confidence intervals around the mean cell number at each time point. Images were converted to cell number estimates every 4 hours. Time of monitoring ranged from 1 week (168 hours) for the untreated control to 4 weeks (672 hours) for the 1000 nM dose. D. Experimental measurements of the t_crit_ for each doxorubicin treatment, legend as in B.

### Integrating estimates of phenotypic composition with longitudinal treatment response data is necessary for identifiable model calibration

To utilize all possible pieces of information available about the treatment response of this experimental system, we sought to develop an integrated model calibration scheme that is capable of integrating information from multimodal data sources. Here, we apply the concept of pareto optimality (27) to reflect the trade-off between goodness-of-fit in each of the two data sources: 1) from longitudinal population data, *N*(*t*), sampled at a high temporal resolution and for a number of doses, and 2) machine learning outputs that estimate the phenotypic composition *ϕ*(*t*) at three distinct time points before and after treatment. For the following dual-objective function, we use a regularization term *λ*, which can vary from 0 to 1 to reflect our varying confidence in the data from each of the measurement sources. Here we use weighted, non-linear, least squares as the simplest possible calibration method

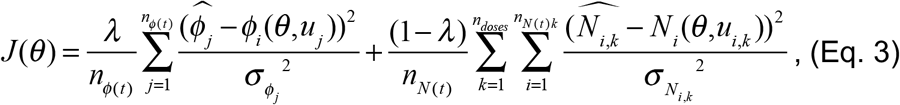

where *n*_*ϕ*(*t*)_ is the number of *ϕ*(*t*) time points, *ϕ_j_* is the experimentally estimated *ϕ* at time point *j, ϕ*(*θ,u_j_*) is the model predicted *ϕ* for a given effective dose *u* at time *j*, *σ*^2^_*ϕj*_ is the variance in the measurement of *ϕ* at time *j*, *n*_*N*(*t*)_ is the number of total *N*(*t*) time points, *n_doses_* is the number of different doses applied, *n*_*N*(*t*)*k*_ is the number of time points in the *k*th dose, *<u>N</u>*_*j,k*_ is the measured number of cells at the *i*th time point for the *k*th dose, *N*(*θ,u*) is the model predicted number of cells at time *i* for the *k*th effective dose, and *σ*^2^_*N*_ is the variance in the measurement of *N* at time *i* for the *k*th dose. The resulting objective function *J*(*θ*), minimizes the sum of the squared error in the *ϕ*(*t*) and *N*(*t*) data compared to the model predicted *ϕ*(*t*) and *N*(*t*). The errors are weighted by the experimentally observed uncertainty in those estimates and normalized by the number of *ϕ*(*t*) and *N*(*t*) data points. The results of this parameter estimation, in terms of weighted error in *N*(*t*) and *ϕ*(*t*) for varying degrees of confidence in each output are shown as the Pareto front set of solutions in Fig 6A. See STAR Methods Model Fitting with Multiple Measurement Sources for a description of how this front was found (Supp Fig S4). The front is centered around *λ*\*=0.5, the regularization term value that equally weights the measurement sources based on the number of data points available from each measurement source. The best fitting parameter set resulting from using the objective function with a *λ*=*λ*\* is denoted as *θ*\* (red dot in Fig 6A) and will be used to evaluate goodness of fit and prediction accuracy going forward.

**Fig 6.**
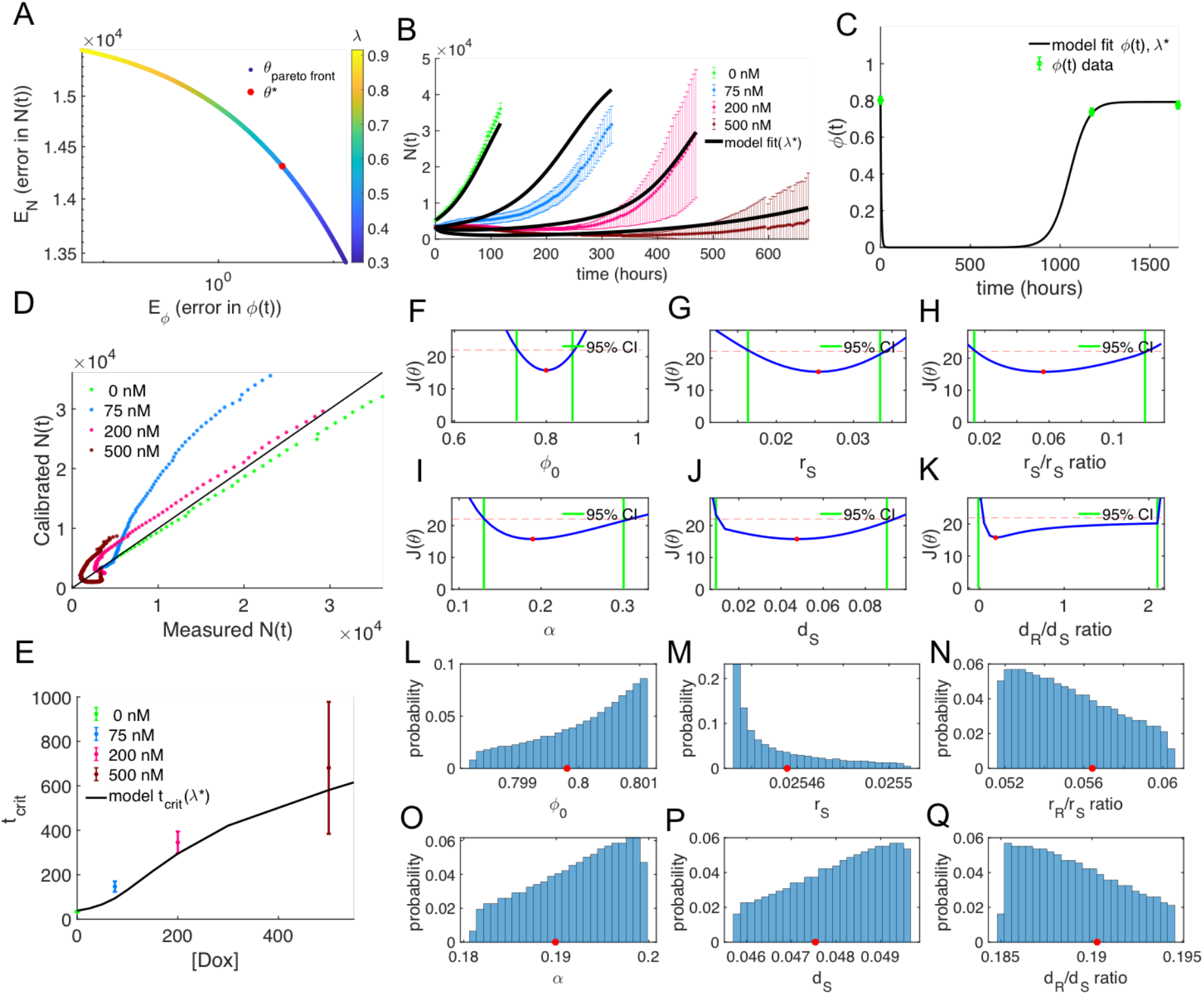
Integrated model calibration to incorporate both measurement sources reveals identifiability of model parameters. A. The parameter sets, *θ*, that fall on the “Pareto front”, reflecting a tradeoff between goodness of fit in *N*(*t*) and *ϕ*(*t*). Each dot represents a parameter set, θ, acquired by varying the regularization term, ***λ***, from 0 to 1 and then filtering solutions (Supp Fig S4) to within a reasonable accuracy in *N*(*t*) and *ϕ*(*t*). The red dot parameter set represents when ***λ*** = ***λ***\*= 0.5 in which the weighting between the measurements sources is given equal weight and normalized based on the number of data points in *N*(*t*) and *ϕ*(*t*) measurements. The Pareto front solutions are found by performing multiple optimizations at different values of ***λ***. Lower values of ***λ*** (blue) indicate improved fit in *N*(*t*) data, whereas higher values of ***λ*** (yellow) indicate improved fit of the *ϕ*(*t*) data. B. Calibration results for longitudinal *N*(*t*) data from the four doses (0, 75, 200, and 500 nM) used for calibration for the parameter set *θ*\* (represented by the red dot in A) C. Calibration results for phenotypic composition (*ϕ*(*t*)) data from the same parameter set *θ*\*, yielding an accuracy in *ϕ*(*t*), measured by the concordance correlation coefficient (CCC) of 0.93 D. Measured cell number *N*(*t*) verses model calibrated cell number, yielding a concordance in *N*(*t*) of CCC = 0.93. E. Critical time from *N*(*t*) data compared to model calibrated critical time for selected parameter set (*θ*\*) in red in (A) (CCC = 0.88). F. Profile likelihood curvature around the initial sensitive cell fraction (*ϕ_0_*) to determine 95% CI on parameter of *ϕ_0_* = 0.80 [0.74, 0.86]. G. Profile likelihood curvature around sensitive cell growth rate (*r_S_*) reveals 95% CI of *r_S_* = 0.026 [0.016, 0.033]. H. Profile likelihood curvature around the ratio of the resistant growth rate to the sensitive cell growth rate reveals a CI of *r_R_*/*r_S_* ratio = 0.056 [0.013, 0.12]. I. Profile likelihood around the drug-induced resistance rate, *α* of *α* = 0.19 [0.13, 0.30]. J. Profile likelihood around the death rate due to drug of the sensitive cell death rate with CI *d_S_* = 0.048 [0.0092, 0.90]. K. Profile likelihood curvature of the ratio of the death rate due to drug of the resistant versus sensitive cell fraction with CI d_R_/d_R_ ratio = 0.19 [-0.014, 2.1]. L. Distribution of Pareto front accepted parameter *ϕ_0_*. M. Distribution of Pareto front accepted parameter *r_S_*. N. Distribution of Pareto front accepted parameter resistant to sensitive growth rate. O. Distribution of Pareto front accepted parameter *α*. P. Distribution of Pareto front accepted parameter *d_S_*. Q. Distribution of Pareto front accepted parameter resistant to sensitive cell death rate.

We use this parameter set, *θ*\*, (red dot in Fig 6A, red dots in Fig 6L-Q) to demonstrate an example of the *N*(*t*) data fit to the model (Fig 6B) and the *ϕ*(*t*) data fit to the model (Fig 6C) with a CCC in *ϕ*(*t*) = 0.93. We demonstrate that the model calibration is fairly accurate at calibrating the *N*(*t*) data (Fig 6D) with a CCC = 0.93, and is able to capture broader changes over the range of doses by properly matching the critical time (*t_crit_*) as a function of dose (Fig 6E) for the four doses the model is calibrated on (CCC = 0.9657). In the model development process, we tested to make sure that each of the parameters was sensitive to the relevant model outputs, in this case the critical time (*t_crit_*) and the phenotypic composition at critical time *ϕ*(*t*=*t_crit_*), for a range of doxorubicin doses. Results from the global sensitivity analysis (See STAR Methods: Sensitivity Analysis of Model Parameters) revealed that all parameters are globally sensitive (i.e. contribute to least 5% of the overall value) in at least one of the model outputs for at least one of the drug doses (Supp Fig. S5), except for the carrying capacities (*K_N_* and *K_ϕ_*) of the two experimental systems. We used this analysis to inform our decision to set the carrying capacities from separate experiments (Supp Fig. S6) and literature (28) and to try to fit all six remaining unknown parameters.

A key goal of this work is not only to fit the model to multiple data sources, but to demonstrate that the use of the information gained from these dispersed data types is critical in enabling the practical identifiability of the six free model parameters in the mechanistic model. The ability to ensure that model parameters are identifiable from data enables us to have confidence in our interpretation of the values of the model parameters to be used for making predictions and ultimately decisions, and thus is essential for eventually translating modeling frameworks like the one presented here to real-world settings. A critical first step is to demonstrate the structural identifiability of the system, which was shown (See STAR Methods: Structural Identifiability of Model Parameters) under the assumption of perfect data. Next, in order to test whether the calibrated parameter set *θ*\* is practically uniquely identifiable from the available data and the objective function (Eq. 3), we utilize the profile likelihood method (29–31). We profiled each parameter independently at a range of values around its best fitting value, *θ*_*i*_*, fitting for all the other parameters, and returning the resulting best possible objective function value (*J*(*θ*)) for the new optimization problem (Fig 6F-K). Parameters that are easily identifiable will result in a tight curvature around the best fitting value, meaning that changing for example the value of the initial sensitive cell fraction (*ϕ_0_*) leads to a large change in the best possible minimized error. Parameters are considered practically identifiable if the curvature of the objective function value crosses above the threshold of the 95% *χ*-squared distribution (32) (red dashed line Fig 6F-K). The parameter value at which this threshold is crossed is considered the upper and lower bound of the 95% confidence interval in the parameter value (green vertical lines Fig 6F-K), providing an estimate of uncertainty in the best fitting parameter value *θ*_*i*_*. The results of this analysis reveal that the six free model parameters are uniquely practically identifiable from the available data. The parameter relationships that result from profiling each individual parameter can be seen in Supp Fig. S7. In contrast, when the test of practical identifiability was repeated for the case in which the calibration was performed on the *N*(*t*) data alone (Supp Fig. S8), and the results revealed a number of non-identifiable parameters in practice (Supp Fig. S9). This analysis demonstrated that the incorporation of the snapshot phenotypic composition data was not only a useful additional piece of information, but essential to making the model calibration and parameter estimates identifiable and ultimately useful.

We further investigated the functionality of our Pareto front set of parameter values by examining the resulting distribution of parameter values that are accepted into the front (Fig 6L-Q). The distributions of parameter values tell us about which parameters, such as the sensitive cell growth rate, *r_S_* (Fig 6L), tend to be very stable regardless of the weighting, whereas other parameters, such as the degree of drug-induced resistance *α*(Fig 6O), are more variable. We observed that the individual parameter values tended to vary systematically with the goodness of fit in *N*(*t*) vs. *ϕ*(*t*) (Supp Fig. S10), however, all of the parameters in the distributions shown in Fig 6L-Q fall within the 95% CI around *θ*\* (Supp Fig. S11). We plot a few examples of the Pareto front solution sets fit to the *N*(*t*) and *ϕ*(*t*) data in Supp Fig. S12, which demonstrates the relatively subtle differences between the fits depending on the weighting of each measurement source.

### Model Validation Using Functional Isolation of “Sensitive” and “Resistant” Cells Predicted from Classifier

Because we rely on the machine learning classifier of cell phenotypes from transcriptomic data, we sought to validate our classifier model experimentally to ensure that cells labeled as “resistant” and “sensitive” were exhibiting these expected phenotypes. Our mathematical model assumes that sensitive cells proliferate more rapidly than resistant cells (i.e. exhibit a higher growth rate) and that resistant cells are capable of improved survival in response to doxorubicin treatment. To test these attributes functionally, we used the COLBERT barcoding system (25) to identify individual lineages from the pre-treatment sample who were labeled as sensitive and resistant based on their changes in lineage abundance, and subsequently isolated them experimentally from the replicate pre-treatment population using the COLBERT recall system (25) (Fig 7A). Once isolated, cells were sorted into single cell clones for functional analysis of growth dynamics and drug sensitivity. Our results confirmed that the cells from the isolated sensitive lineage grow more quickly than the isolated resistant lineage (Fig 7B), with overall growth rates of *g_S_* = 0.011 and *g_R_* = 0.005 per hour respectively (Supp Fig S13). Drug sensitivity was assessed by dosing cells at 400 nM and 2.5 μM for 48 hours and immediately quantifying cell viability via a live-dead assay. The resistant lineage had higher percent viability at both doxorubicin concentrations, with a statistically significant difference in viability at the higher dose (Fig 7C).

**Fig 7.**
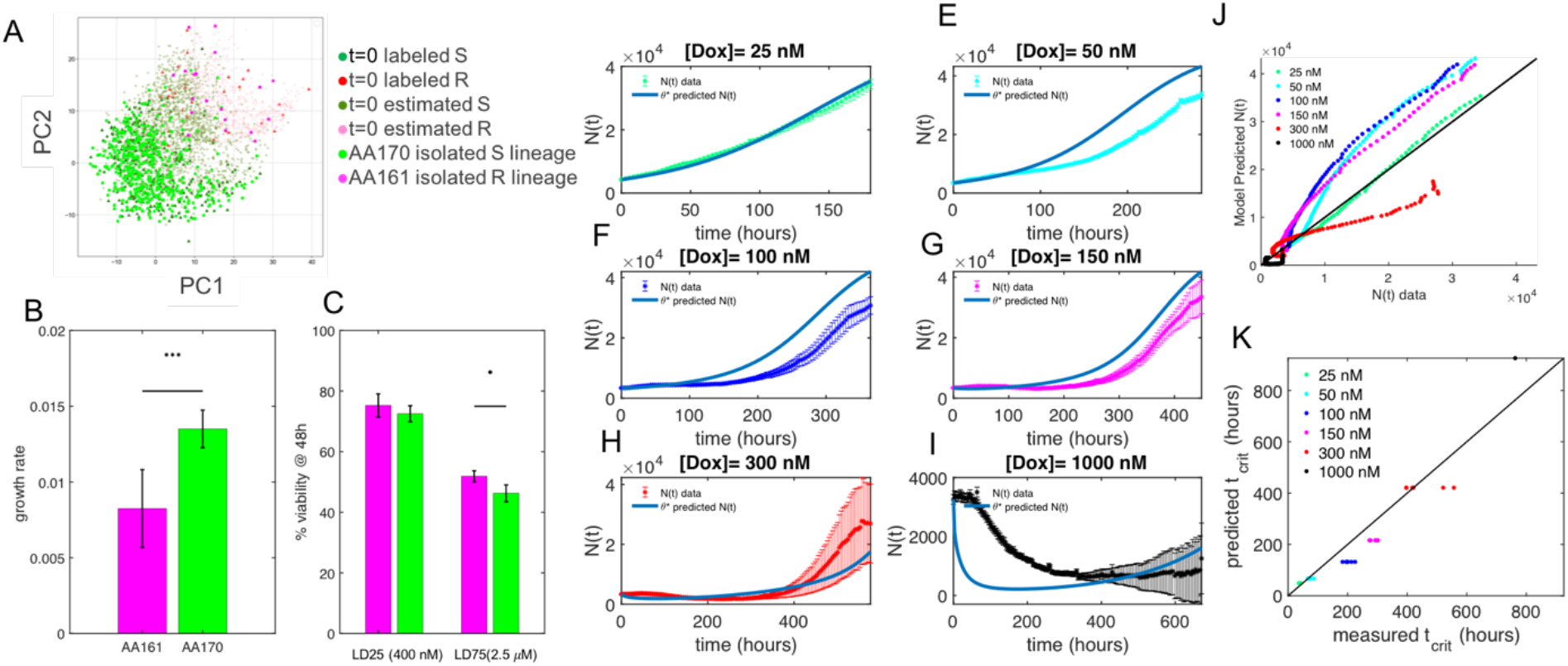
Combined Model Validation *via* Lineage Isolation and Prediction of Treatment Response. A. Projection of classified sensitive and resistant cells at the pre-treatment time point into principal component space, with cells from an isolated sensitive lineage (AA170) in bright green, and an isolated resistant lineage (AA161) in hot pink B. Growth rates of the 12 replicate wells of each isolated lineage reveal that the resistant lineage grows significantly more slowly than the sensitive lineage (p = 2.7e-6), as is predicted from the model parameters where *r_S_* > *r_R_*. C. Functional testing of the drug sensitivity of each lineage indicates that the cells from resistant lineage (AA161, pink) have a higher resistance, measured by cell viability at 48 hours, at both 400 nM and 2.5 μM doses of doxorubicin, with p-values of p = 0.1942 and p = 0.0023, respectively. D. Prediction of treatment response at 25 nM, E. 50 nM, F. 100 nM, G. 150 nM, H. 300 nM, and I. 1000 nM from *θ*\* (red dot in Fig 6A). The mean measured cell number in time and 95% confidence interval from six replicate wells are shown for each treatment response. J. Scatterplot of model predicted *N*(*t*) versus experimental *N*(*t*) data for all 6 new treatment conditions with an overall CCC = 0.89. K. Scatterplot of model predicted critical time from selected parameter set versus experimentally measured critical time, indicating that although we might not be able to precisely predict the exact trajectories of cell number in time for each dose perfectly, we can globally capture the critical time (*t_crit_*) for a range of doxorubicin concentrations, despite our model not being trained on these concentrations, with an overall CCC between model predicted critical time and observed critical times of CCC = 0.92.

### Multimodal Data Sources can be Leveraged to Predict Response Dynamics to New Drug Concentration

A key advantage of leveraging multimodal data sources for parameter estimation is that we can uniquely identify the model parameters and use them to make predictions about the response dynamics to new treatment regimens. We validate the model predictions, obtained from running the model forward with parameter set θ* with input effective doses described in Fig 5B for the six remaining pulse treatment of doxorubicin that were not used to train the model. The model predictions compared to the experimental measurements are shown for doses of 25 nM (Fig 7D), 50 nM (Fig 7E), 100 nM (Fig 7F), 150 nM (Fig 7G), 300 nM (Fig 7H) and 1000 nM (Fig 7I). We evaluated the accuracy in all the model predictions over all six unobserved doses and see that we are able to predict the treatment response with reasonable accuracy (Fig 7J) with an overall CCC of 0.89 for each model predicted and measured cell number (*N*(*t*)) in time. When we repeat this calibration, removing that phenotypic composition data (by setting *λ* = 0) we get an overall predictive accuracy of CCC=0.79, indicating the improvement in predictive capabilities with insight of the phenotypic dynamics. While the individual trajectories may not precisely match the data at the 4-hour intervals measured here, they are able to predict the global behavior, *via* predicting the critical time as a function of doxorubicin, very well (Fig 7K) with a CCC of 0.92. These results demonstrate the flexibility and predictive capability of this modeling framework, demonstrating its utility in predicting the critical behavior needed to guide optimal-treatment decision making.

## Discussion

Recent technological advances have enabled unprecedented, high-throughput single-cell molecular level insight of intratumor heterogeneity(33,34). The ability to precisely quantify intratumor heterogeneity (1), and illuminate key subpopulations involved in response to treatment (11), has the potential to improve both prognostic and therapeutics for cancer treatment. These genomic and transcriptomic data sets can direct the choice of specific cancer drugs and illuminate novel resistance pathways, as well as provide a prognostic marker for patients who receive it. Simultaneously, the role of mathematical modeling in oncology has been widely recognized (35) and utilized to improve both our understanding of the dynamic mechanisms of drug response (12,36,37) as well as to develop approaches to guide the design of adaptive patient-specific treatment plans (14,19,20,38,39). However, connecting the wealth of “omics” data at the molecular level with temporal dynamics used to calibrate mathematical models for adaptive therapies remains a major challenge in the field.

Recognizing the critical roles of heterogeneity in cancer dynamics, mathematical models of tumor progression often include distinct subpopulations, such as cancer stem cells (14,40,41), or drug resistant and sensitive subpopulations (17,18,21,42). However, despite these models being calibrated to observed experimental or clinical data, the underlying phenotypic composition that these model calibrations suggest cannot easily be validated, since the degree of resistance or stemness of a cancer cell population in time is not easily measured longitudinally *via* a single biomarker. The majority of these modeling endeavors utilize a single measurement source for longitudinal data acquisition and subsequent model calibration. A few studies utilizing multimodal imaging modalities have harnessed the ability to quantify different aspects of tumor composition—such as vasculature, necrosis, and cellularity, to develop an integrated model calibration of multiple tumor system components (43,44). However, this integrated, multimodal approach has not explicitly included inference of the composition of heterogeneous subpopulations taken from separate “omics” datasets for direct model calibration.

Here, we introduce an experimental-computational framework for utilizing multimodal data sets when parametrizing a mechanistic model of drug resistance dynamics in response to treatment in cancer. We demonstrate the applicability of this framework when applied to clonally-resolved scRNA-seq data combined with longitudinal treatment response data from a cancer cell line and assess the ability of the model to predict treatment response dynamics. To this end, we developed a machine learning classifier built upon clonal abundance quantification which estimates the class identity of an individual cell based on its transcriptome. The machine learning outputs classified cell states and were used to assign values to the state variables in the mechanistic model: the number of cells in the sensitive or resistant phenotypic state at each time point. We combined these estimates of phenotypic composition with population-level treatment response data to calibrate a mechanistic model of drug-resistance dynamics. We validated our machine learning classifier by isolating cells from lineages labeled as sensitive or resistant and testing them functionally. We showed that the presence of multiple measurement sources of data allows for the practical identifiability of the model parameters, which are then used to accurately predict the effect of new drug treatments on the cell population.

The power of mathematical models in oncology, especially those calibrated to real data, is that we can both use them to learn about the underlying mechanisms of the system behavior, and we can harness that knowledge to inform future decision making in an experimental or clinical setting (45,46). Greene *et al*. (17) demonstrate that knowledge of the parameters of the model presented here (Eq.1) can be used to drive optimal treatment protocol decisions; in particular they can help determine (for example) whether pulsed or constant treatment is preferred for a specific patient. The applicability of optimal control theory as it applies to cancer treatments relies on the ability to identify model parameters from data. While it has been shown that the model parameters presented in this paper can be identified from just the bulk population dynamics in theory (16), in practice the number of experiments needed to test the conditions is quite difficult if the output is the bulk population dynamics alone. However, the identifiability problem becomes significantly easier if the knowledge of the underlying phenotypic composition is also plausible (See STAR Methods Identifiability of Model Parameters). In this work, we leverage high-throughput “omics” data sets, taken at just a few snapshots of time, to estimate the phenotypic composition and demonstrate the improvement in identifiability of model parameters from including this data alongside longitudinal data.

High-throughput single cell transcriptomics or other types of high throughput snapshot data can give an abundance of information about the heterogeneity and potential mechanisms of resistance of cell populations (11,47). However, the ability to use this information beyond hypothesis generation (12), but to actually inform model calibrations, is still lacking. In this work, we attempted to overcome the problem of practical identifiability of model parameters from observed data by demonstrating how to explicitly integrate snapshot data about the relevant cell subpopulations into a model calibration. We argue that the ability to integrate information from snapshot data with temporal data is essential for the potential for the proposed mathematical oncology models to be practically useful, as these models should not “throw away” information but should instead be able to take into account explicitly as much available data as possible.

The functional characterization of single cells via changes in lineage abundance post-treatment enabled us to identify novel mechanistic insights into which pathways and interactions are critical for surviving treatment response. While clustering of cells by their transcriptomes can enable identification of novel cell states, these cell states are not necessarily relevant to drug-tolerance. Once can see this quite simply in scRNA-seq pipelines as failure to remove cell cycle genes from the analysis reveals that cells will often cluster by cell cycle state (48). While states of the cell cycle may be important for certain applications, they are often regressed out. However, we cannot regress out other unknown phenotypic subpopulations, and thus these are what can emerge from unsupervised clustering algorithms. While these can provide novel insight about population structure, they may not be what is relevant to driving changes in treatment response behavior. Thus, the ability to read-out lineage identities represents a novel functional component that enables us to zoom in at the right phenotypic state-space relevant to our question-what cells are more drug resistant and which are more drug sensitive, and what is driving these changes? Because we used principal component analysis to build a classifier to separate the sensitive and resistant cells, we can look at the differences in gene expression patterns between the groups of cells we identified and propose potential novel interactions and new biomarkers. For example, our analysis reveals TOP2A, NEAT1, and UBE2S as delineators between sensitive and resistant cells. This knowledge can provide the basis for future work investigating the role of these genes and their related pathways in drug-response.

While scRNA-seq has limitations in the clinical setting due to its high cost, in experimental settings barcode labeling fits rather flexibly into existing scRNA-seq workflows and can add a critical functional component to the phenotypic read-out, as we display in this work. In the clinical setting, other types of approaches to learn more about cancer cell composition are being employed in the era of precision medicine. From radiomics to genomics, it is becoming increasingly common for patients to have access to high-throughput measurements, or at least some insight into their mutational burden at certain time points. This information may be integrated into the clinical or tumor board’s decision-making process (49).

We suggest that the approach here could be modified for the available types of snapshot data in different experimental or clinical settings, where it is expected that only sparse data in terms of tumor composition and longitudinal dynamics will be available. This could potentially be overcome in two ways: 1) simplifying the model structure to reduce the number of model parameters, or 2) setting some parameter values to those obtained from the literature, and only allowing a few (key) parameters to be patient-specific, as is performed in (14). We anticipate that this integrated approach can be applied flexibly to incorporate and integrate snapshot data about population composition with longitudinal bulk population dynamics. While transcriptomic and longitudinal data have been used together in a number of studies, this is the first work to our knowledge that allows for explicit parameter estimation using multimodal measurement sources of varying time resolutions and enables flexible implementation depending on the degree of confidence in each data source. The synergy of machine learning with mechanistic modeling integrates multimodal datasets and opens up new approaches to describe, predict, and ultimately optimize treatment response in cancer.

## STAR Methods

### Key Resources Table

See Attached Template

### Experimental Model and Subject Details

### Cell culture

The human breast cancer cell line MDA-MB-231(ATCC) was used throughout this study. Cells were maintained in Dulbecco’s Modified Eagle Medium (Gibco) and supplemented with 1% Penicillin-Streptomycin (Gibco) and 10% fetal bovine serum (Gibco) under standard culture conditions (5% CO_2_, 37°C).

A subline of the MDA-MB-231 breast cancer cell line was engineered to constitutively express EGFP (enhanced green fluorescent protein) with a nuclear localization signal (NLS). Genomic integration of the EGFP expression cassette was accomplished through the Sleeping Beauty transposon system (50). The EGFP-NLS sequence was obtained as a gBlock from IDT and cloned into the optimized sleeping beauty transfer vector containing the EGFP-NLS expression cassette and the pCMV(CAT)T7-SB100 plasmid containing the Sleeping Beauty transposase was co-transfected into a MDA-MB-231 cell population using Lipofectamine 2000. mCMV(CAT)T7-SB100 was a gift from Zsuzsanna Izsvak (Addgene plasmid #34879) (51). GFP+ cells were collected by fluorescence activated cell sorting. MDA-MB-231 cells are maintained in DMEM (Gibco), 10% fetal bovine serum (Gibco) and 200 μg/mL G418 (Caisson Labs). Cells were seeded into the center 60 wells of a 96 well plate (Trueline) at about 2000 cells per well. During the monitoring and treatment, plates were kept in the Incucyte Zoom, a combined incubator and time-lapsed microscope. Cells were fed fresh media every 2-3 days for up to 5 weeks. HEK293T cells were cultured in DMEM with GlutaMAX supplemented with 10% FBS, 4.5 g/L D-glucose, 110 mg/L sodium pyruvate, streptomycin (100ug/mL) and penicillin (100 units/mL).

### Longitudinal treatment response data

The EGFP-labeled subline of MDA-MB-231 breast cancer cells were used for longitudinal treatment response. Cells were passaged into the center 60 wells of 96 well plates at a density of about 2000 cells per well. Two days later, cells were treated with a 24 hour pulse-treatment of doxorubicin at concentrations ranging from 0-1000 nM, with 6 replicates of each dose. Dosed media was applied to cells and treatment response was monitored using the Incucyte. After 24 hours, the dosed media was replaced with normal media and monitoring continued. Cells were fed fresh media every 2-3 days for the duration of the monitoring period (up to 5 weeks).

### Integration, expression, and capture of COLBERT barcodes

#### Lentiviral Assembly

Lentiviral assembly was performed using the Lenti-Pac HIV Expression Packaging Kit (GeneCopeia). Two days prior to lentiviral transfection 1.5 million HEK293T cells were plated in a 10 cm tissue culture dish. Forty eight hours after plating, cells were 70-80% confluent and transfected with 9 μL of Lipofectamine 2000 (Thermo Fisher #11668027), 1.5 μg per well of PsPax2 (Addgene #12260), 0.4 μg/well of VSV-G (Addgene #8454), and 2.5 μg of Lenti-Pac HIV mix (GeneCopoeia). Media was replaced 24 hours post transfection with 10 mL DMEM supplemented with 5% heat inactivated FBS and 20 μL TiterBoost (GeneCopoeia) reagent. Media containing viral particles was collected at 48 and 72 hours post transfection, centrifuged at 500g for 5 minutes, and filtered through a 45 μm poly(ether sulfone) (PES) low protein binding filter. Filtered supernatant was stored at −80 °C in aliquots for later use.

#### Barcode Labeling

MDA-MB-231 cells were transduced with the Cropseq-BFP-WPRE-TS-hU6-N20 lentivirus in growth media with 1 μg/mL polybrene. After 48 hours of incubation, 1000 BFP+ cells were isolated by FACS to establish a population with initial diversity of ~1000 unique barcodes. To reduce the likelihood that two viral particles enter a single cell, the lentiviral transduction multiplicity of infection was kept below 0.1.

#### Drug Treatment of Barcoded Cells for scRNAseq and Recovery

Barcode labeled MDA-MB-231 cells (5 replicate wells) were treated with doxorubicin (550 nM) for 48 hours in growth media, washed and replaced with fresh growth media. Surviving cells were maintained in growth media and passaged up serially from 0.1 x 10^6^ to 20 x 10^6^ cells.

#### scRNA-seq

Cryopreserved samples from drug-naïve and two samples of doxorubicin-treated cells frozen at 7 and 10 weeks post-treatment were harvested, sorted by FACS to collect the BFP+ population. Cells were loaded into wells of a Chromium A Chip, and libraries were prepared using the 10XGenomics 3’ Single Cell Gene Expression (v2) protocol. Paired end (PE) sequencing of the libraries was conducted using a NovaSeq 6000 with an S1 chip (100 cycles) according to the manufacturer’s instructions (Illumina).

#### Plasmid Assembly for Isolation of Lineages

After selecting the lineages of interest for isolation, an array of barcodes was assembled as described in (25). Briefly, oligonucleotide pairs for the barcode of interest were ordered with specific overlapping sequences to both direct assembly of barcode array and integration into the plasmid for isolation. The barcode arrays were ligated, and gel purified to proceed with only a fully assembled array in cloning. The fully assembled barcode array was cloned into the BbsI site with standard restriction digest cloning. This double stranded barcode array was inserted into a plasmid backbone upstream of a minimal core promotor (miniCMV) and sfGFP to generate the Recall plasmid. This was repeated with individual barcodes of interest.

#### Recall of Isolated Sensitive and Resistant Clones by COLBERT

Barcoded MDA-MB-231 cells were seeded in 6 well plates and transfected using Lipofectamine 3000 (ThermoFisher) with 225 ng dCas9-VPR-Slim and 275 ng Recall Plasmid per well. Forty eight hours after transfection, GFP+ cells were single cell sorted by FACS into a 96 well plate and spun for 1 minute at 1000g. Sorted cells were expanded until 80% confluency and passaged into a single well of a 48 well plate. Upon first passage following sort, 1/6 of the cells or ~5000 live cells were resuspended in a PCR reaction mix to confirm lineage identity through PCR amplification and subsequent Sanger sequencing of barcode region.

#### Alignment to Reference Genome

The GTF file included with cellranger’s GRCh38 3.0.0 reference was modified to create a “pre-mRNA” GTF file so that pre-mRNAs would be included as counts in the later analysis. Cellranger’s (v3.0.2) *mkref* command was then used to create a pre-mRNA reference from the GTF file and a genome FASTA file from the GRCh38 3.0.0 reference. FASTQ files of the scRNA-seq libraries were then aligned to the new pre-mRNA reference using the *cellranger count* command, producing gene expression matrices. The matrices for the different samples were concatenated into a single matrix using the *cellranger aggr* command with normalization turned off, so that the raw counts would remain unchanged at this point.

#### Filtering and Normalization

The filtered matrices produced by cellranger were loaded into scanpy (v1.4.4)(52). Cells were annotated by sample and lineage membership. Only cells meeting the following requirements were retained for further analysis: (a) a minimum of 10000 and maximum of 80000 transcript counts, (b) a maximum of 20% of counts attributed to mitochondrial genes, and (c) a minimum of 3000 genes detected. Genes detected in fewer than 20 cells were removed. Normalization was conducted based on the recommendations from multiple studies that compared several normalization techniques against each other(48,53,54). In brief, three steps were performed: (a) preliminary clustering of cells by constructing a nearest network graph and using scanpy’s implementation of Leiden community detection(55), (b) calculating size factors using the R package scran(56), and (c) dividing counts by the respective size factor assigned to each cell. Normalized counts were then transformed by adding a pseudocount of 1 and taking the natural log.

#### Regressing Out Cell Cycle Expression Signatures

Using a list of genes known to be associated with different cell cycle phases (57), cells were assigned S-phase and G2M-phase scores. The difference between the G2M and S phase scores were regressed out using scanpy’s *regress_out* function.

### Quantification and Statistical Analysis

#### Machine Learning of Cell Phenotypes

The machine learning classifier of sensitive and resistant cell phenotypes was built from the normalized, pre-processed single cell gene expression matrix with lineage identities from the pre-treatment time point only. For the cells in the pre-treatment sample, the lineage abundance at the pre-treatment time point (proportion of cells in each lineage) was calculated and compared to the lineage abundance at the combined post-treatment time points. If the lineage was not observed in the post-treatment time points, the lineage abundance post-treatment was assigned a zero. The change in lineage abundance (% post -% pre) was found for each lineage in the pre-treatment time point (See Fig. 3A). Based on this change in lineage abundance distribution, the pronounced tails of the distribution were used for classification. Cells whose lineage abundance change was positive, i.e. the lineage abundance increased post-treatment, were labeled as resistant in the pre-treatment time point. Cells whose lineage abundance change decreased by more than 5% were labeled as sensitive in the pre-treatment time point. This resulted in 815 cells and their corresponding 20,645 normalized gene expression levels being used to form the training set gene-cell matrix containing a cell’s gene expression vector and corresponding identity (with a 0 being sensitive and a 1 being resistant). This gene-cell matrix was then used to build a classifier capable of predicting the identity of new cells based on an individual gene expression vector.

A principal component classifier was built using the methods for eigendigit classification proposed in (58) for performing face recognition. In short, we perform principal component analysis of the gene-cell matrix from the labeled sensitive and resistant cells only. Each labeled cell’s gene expression vector is projected into principal component space, made up of the gene loadings of each of the 20,645 genes and the optimal number of principal components. To classify new cells based on their gene expression and corresponding projection into PC space, the k closest cells to the new cell in the labeled class are found using a Euclidian distance metric. If the new cell’s k neighbors give the cell a probability of being resistant of greater than a certain threshold, than that cell is classified as resistant. If it is less than the cut-off threshold, it is classified as sensitive. This process is repeated for all unclassified cells from the remaining pretreatment time points and all of the post-treatment time points. All calculations of principle component coordinates and *knn*-probabilities were found using python’s scanpy package. The number of components, neighbors, and threshold probability were optimized via coordinate optimization described below.

#### PCA Classifier Hyperparameter Optimization

Using the labeled sensitive and resistant cells from the pre-treatment time point, 5-fold CV was used to split the cells into evenly class-balanced groups of training and testing data sets. Coordinate optimization was then used to iteratively find the optimal number of both nearest neighbors (*k*) and number of principal components (*n*) for correctly identifying the class of each cell. Coordinate optimization works by essentially iteratively optimizing the two variables of interest, here k and n, until they no longer change values. In this case, we first set the number of principal components to a single value and iterated through a range of nearest neighbors to find the number which gave the highest mean AUC (area under the curve) over all 5 folds of cross validation (Supp Fig. S1A). Once the optimal number of neighbors was found for that number of principal components, the number of neighbors was set to that value and the optimal number of principal components was varied over a range of values, and again the highest mean AUC over all 5 folds of cross validation was found (Supp Fig S1B). Then we set the number of neighbors to this value, and repeated the search for the optimal number of principal components. This process was repeated until the optimal number of neighbors and number of principal components no longer changed with each iteration. Using the optimal hyperparameters, we projected all of the labeled cells into the full classifier model and found the ROC curve for different probability thresholds for classifying cells as sensitive or resistant. While many appeared to be reasonable, we chose a threshold value of P(resistant)= 0.2 as our cut-off for calling a cell resistant, as this generated a realistic proportion of cells in each class at the pre-treatment time point.

#### Model of Drug Resistance Dynamics

The mathematical model of drug-induced resistance, in which treatment exposure directly induced phenotypic transitions into the resistant cell state, was introduced in (17). Their original model described sensitive cells (*S*) and resistant cells (*R*) independently growing according to logistic growth and independently dying due to drug treatment (*u*(*t*)) *via* a log-kill hypothesis. The model includes an explicit role for the transition of sensitive cells into resistant cells via a rate of drug-induced resistance (*α*) which is modeled as a linear function of treatment *u*(*t*). Additionally, their full model included additional terms of spontaneous, treatment-independent resistance (*ε*) proportional to the number of sensitive cells present, as well as a resensitization term (*γ*) describing treatment-independent transition from the resistant to the sensitive cell state.

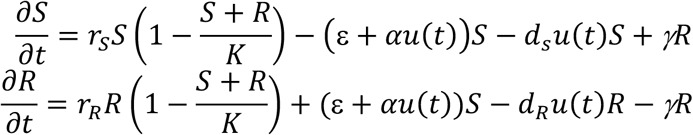

In order to have the best possible chance of identifying these model parameters from data, we simplified the original model. We assume that the treatment-independent transition into the resistant state (*ε*) and the resensitization (*γ*) are negligible, yielding the following system of equations.

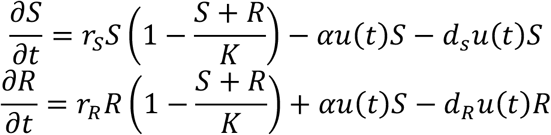

Where *r_S_* and *r_R_* are the sensitive and resistant subpopulation growth rates and *d_S_* and *d_R_* are the sensitive and resistant subpopulation death rates, assumed to be linearly proportional to the effective dose (*u*(*t*)). We assume that the sensitive cells grow faster than the resistant cells so that r_s_ > r_r_, as is consistent with the mechanism of action of cytotoxic therapies targeting rapidly proliferating cells (17,59). We assume *d_S_* > *d_R_* as sensitive cells should die more quickly in response to drug than resistant cells, by definition. We modeled the effect of the pulse-treatments as single pulses of *u*(*t*) whose maximum is given by the concentration of doxorubicin and whose effectiveness in time decays exponentially.

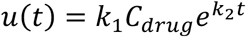

The constants *k_1_* and *k_2_* were chosen so that *u*(*t*) is scaled between 0 and 5 and so that the effective dose decays over a time scale consistent with experimental observations of doxorubicin fluorescent dynamics *in vitro* (22,23). Numerical simulations of the forward model for a given treatment regimen were implemented in MATLAB using the backward Euler method.

To evaluate and compare the effect of treatment regimens on the cell population, we utilized an unbiased time-to-event metric proposed for use in evaluating treatment benefit in clinical trials (24), (there called TTB120 time to reach *1.2*N_0_*), and here which we call critical time or *t_crit_*, defined as the time to reach *2*N_0_*. The longer the *t_crit_*, the longer the “tumor burden” is held below this threshold, and therefore the more effective the treatment regimen. This critical time can be simulated for a given *u*(*t*) in our model and can also be measured experimentally for most doses administered. We therefore use this output, as well as the phenotypic composition at *t_crit_*(*ϕ_s_*(*t*=*t_crit_*)), as outputs for performing sensitivity analysis to assess the effect of parameters on the observed drug response.

#### Sensitivity Analysis of Model Parameters

As part of the model development process, we performed a sensitivity analysis to assess the effect of individual model parameters on the model output. Although there are a number of choices to use for model outputs, we chose to capture the broad drug response of the population using the critical time (*t_crit_*), and the phenotypic composition *ϕ*(*t*=*t_crit_*) at that time, as we expect these are two outputs we would feasibly observe in an experimental setting, as the time to population rebound and the phenotype observable via scRNAseq or some other phenotypic characterization. We first performed a global sensitivity analysis on the set of parameter bounds that were well outside the parameter ranges of the calibrated parameters and their associated errors. The results of the sensitivity analysis will reveal the most important parameters of the system, causing the greatest variation in outputs. This exercise should identify any model parameters that the model is insensitive to, and therefore may present opportunities to simplify the model to capture the same dynamics while reducing uncertainty by eliminating the number of free parameters to be fit. A Sobol’s global sensitivity method is applied, which is a method that utilizes the analysis of variance (ANOVA) decomposition to define its sensitivity indices(60). As a global method, random sampling is performed twice over the parameter space of the eight parameters (six free, two carrying capacities), with the number of parameters by *N* simulations matrices denoted by X and Z. The bounds of the global sensitivity analysis were chosen to be well outside of the 95% confidence intervals around each best fitting parameter from the profile likelihood analysis. The total effects are then calculated using the following:

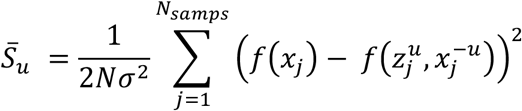

Where *σ*^2^ is the variance of the outputs from the first set of N random samples computed from evaluating over all *x_j_* in *X*, and the function evaluations of *f*(*x_j_*) and 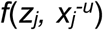 are the outputs (*t_crit_* or *ϕ*(*t*=*t_crit_*)) of the model at parameter values *x_j_* compared to the function evaluated at parameter values *z_j_* for one parameter, and *x_j_* for all the remaining parameters. The total effects were calculated for each parameter value for outputs of both critical time (*t_crit_*) and phenotypic composition (*ϕ*(*t*=*t_crit_*)) for four doses ranging from 0 to 500 nM. Large sensitivity indices between parameters and model outputs characteristics indicate that small changes in the parameter values will result in large variations in the output behavior. For this investigation, to ensure the convergences of the indices, a base simulation size of *N*=5000 is chosen, resulting in (5000 x 2 x 4 doses x 2 outputs x 8 parameters=640,000) simulations to generate the indices. For this study, only the total effects of the model outputs of *t_crit_* and *ϕ*(*t*=*t_crit_*) are reported. Specifically, the critical time and phenotypic composition at critical time is recorded for each random simulation and each dose, and per the Sobol method, the total effects indices derived from the variances of these outputs is calculated, which account for variations in individual parameters as well as additional effects resulting from the combined variation of parameters. A sensitivity cut-off of 0.05 is used, indicating parameters that cause less than 5% of the total variation of that model output.

To perform a local sensitivity analysis, we varied each parameter independently from a single parameter set chosen from the set of Pareto optimal sets. To perturb each parameter, we chose a high parameter value of two times its optimal value, and a low parameter value of half its optimal value. We used these high and low parameter values, holding all other parameters constant, and ran the forward model and recorded the response over a range of doxorubicin doses from 0-500 nM, for both the effect in critical time (*t_crit_*) and phenotypic composition at critical time (*ϕ*(*t*=*t_crit_*)). For each independent parameter perturbation, we computed a high and low sensitivity score for the the ith parameter, for the two model outputs (*t_crit_* or *ϕ*(*t*=*t_crit_*)) as:

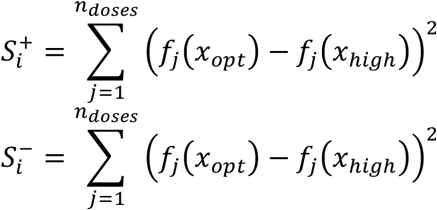

Which is the sum-squared difference between the output values (*t_crit_* or *ϕ*(*t*=*t_crit_*)) for each jth dose in the range of doses, for both the high and low parameter sets, for each ith parameter. The sum of the high and low sensitivity scores for each parameter were than ranked for the two outputs of *t_crit_* and (*ϕ*(*t*=*t_crit_*)). This analysis reveals the most important parameter in driving changes in output behavior of the model locally around the best fitting parameters.

#### Model Fitting with Multiple Measurement Sources

To perform model fitting, we used two sources of measurement data: cell number in time (*N*(*t*)) in response to the pulsed doxorubicin treatments, and estimates of the phenotypic composition, *ϕ*(*t*), at three time points total (before and two post-treatment). The data were collected in two separate experimental settings, with two different carrying capacities, which we refer to as *K_N_* and *K_ϕ_*. The longitudinal cell number data was recorded in 96 well plates, resulting in a different carrying capacity than the lineage-traced single cell RNA sequencing experiment in which the population was expanded out to a 15 cm dish due to the need for large cell numbers for running on the 10x Genomics system. The carrying capacity of the longitudinal data, *K_N_*, was found by fitting the untreated control to a logistic growth model and allowing both the effective growth rate of the total population (*g_eff_*) and *K_N_* to be fit to the data (See Supp Fig. S11).

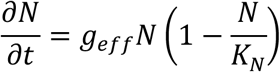

We set this carrying capacity in the model going forward for fitting the longitudinal data. For the carrying capacity of the single cell RNA sequencing experiment, *K_ϕ_*, we used Thermo-Fisher published “Useful Numbers for Cell Culture” as an estimate(28), where the manufacturer cites the number of cells at confluency of 20 million cells. Going forward, we fit the remaining 6 parameters of *θ*=[*ϕ_0_, r_S_, r_S_/r_R_ ratio, α, d_S_, d_R_/d_S_ ratio*] where these represent: the initial fraction of sensitive cells prior to treatment, the sensitive cell growth rate, the ratio of the resistant to sensitive cell growth rate, the rate of drug-induced resistance, the sensitive cell death rate, the ratio of resistant to sensitive cell death rates, respectively. All six parameters were found to be globally sensitive in one or more of the treatment conditions when looking at either *t_crit_* or *ϕ*(*t*=*t_crit_*), and so we decided it was reasonable to try to fit them all from the observed data. We note *r_S_/r_R_* ratio and *d_S_/d_R_* ratio are used for ease of parameter estimation. Since we assumed that *r_r_<r_s_* and *d_r_<d_s_*, we can search the ratio, *r_S_/r_R_* ratio = *r_R_/r_S_* and *d_S_/d_R_* ratio = *d_R_/d_S_*, between 0 and 1 when performing parameter estimation.

To estimate the model parameters *θ*, we used both measurement sources *N*(*t*) and *ϕ*(*t*) and a regularization term, *λ*, which is allowed to vary between 0 and 1 to reflect varying degrees of confidence in the two measurements sources. The data were fitted using a weighted-sum-of-squares-residual function described below:

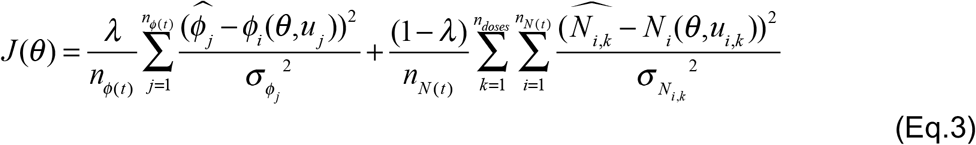

For the *N*(*t*) data, the uncertainty in the data (*σ*^2^_*N*_) at each time point was quantified using the standard deviation of the cell number over the six replicate wells. For the uncertainty in the *ϕ*(*t*) estimates, we compute the Bernoulli sample variance of

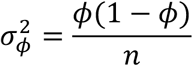

where n is the number of Bernoulli samples (which here is the number of cells in the data set) at each of the three time points. However, this leads to an underestimate in the uncertainty in the *ϕ*(*t*) estimates, which depend significantly on where the threshold is chosen. For this reason, we added an uncertainty term of technical noise *σ_tech_* = 0.01 to this estimate. In reality, the magnitude of the uncertainty in the *ϕ*(*t*) is not necessarily known, and the introduction of the regularization term, *λ*, in practice allows us to vary the degree of certainty we have in each measurement source relative to the other.

The key feature of the introduction of the regularization term *λ* means that we can tune the joint objective function to favor minimizing error in *N*(*t*) and *ϕ*(*t*). In other work in the biomedical field using multi-objective function optimizations, the number of data points from each measurement source is typically similar, as most data is acquired longitudinally (43). However, in this case we have significantly higher time and dose resolution in our *N*(*t*) data (472 data points) compared to our *ϕ*(*t*) data (3 data points), and thus chose to include normalization terms in our objective function (Eq. 3) to account for the different resolutions of the data *N*(*t*) and *ϕ*(*t*) data. Because the data come from distinct measurement sources, the robust quantification of comparative uncertainty is not known *a priori*, as we do not intuitively know whether or not the *ϕ*(*t*) estimates from scRNA-seq are inherently more or less reliable than the longitudinal population size data. We expect this problem to be present for any measurements taken from different measurement sources. Thus, we introduce the regularization term *λ*, which enables tuning of the certainty in favor of one measurement source over the other. We observe a trade-off in goodness of fit where if we assign a high value to *λ*, very close to 1, this puts more confidence on our *ϕ*(*t*), and if we assign a lower value to *λ*, this puts more confidence in our *N*(*t*) data, and favors minimizing the error in that fit.

We first perform parameter estimation using weighting normalized only by the number of data points, with a value of *λ* = 0.5 which we call *λ*\*. We use a multistart search algorithm where we randomly initialize the guess of the initial parameter vector over a range of reasonable parameter space for 100 initial parameter sets. We use the *fminsearch* function in MATLAB to search for a set of parameters, *θ*, that minimized *J*(*θ*) for *λ* = *λ*\* for each initial guess. We then select the parameter set that produces the lowest objective function value, *J*(*θ*). The results of this optimization are presented in order to show an example of a single calibrated parameter set compared to the observed data, as well as to test the identifiability of those parameter values, which we call *θ*\*. This set of parameter values was also used for the local sensitivity analysis.

In order to allow for flexibility and generalizability of the approach for multimodal data sets, we sought to find more than a single optimal parameter set, but a “front” of solutions that could take into consideration the potentially varying degrees of confidence in the two types of measurement sources. We pulled from the field of economics to introduce a concept known as Pareto optimality (27), in which our set of Pareto optimum parameter sets reflects solutions in which an improvement in the fit to *N*(*t*) leads to a trade-off resulting in a worse fit in *ϕ*(*t*). To find the Pareto front set of solutions, we varied the regularization term *λ* from one which only considers the *N*(*t*) data (*λ*=0), to one which only considers the weighs the *ϕ*(*t*) data (*λ=1*). We generated a vector of 1000 ordered *λ* values and iterated through 1000 optimizations at each value of *λ*. We used the concept of homotopy continuation (61) to initialize the guess for each optimization as the best fitting parameter set θ from the previous iteration. For each optimization, we recorded all of the parameter values, the sum-of-squares error in *N*(*t*), the sum-of-squares error in *ϕ*(*t*) the CCC in *N*(*t*), and the CCC in *ϕ*(*t*). The results of the initial optimization are shown in Supp. Fig. S4A, colored by their *λ* value. Next, we filtered the parameter sets, by only keeping those whose parameter values led to a CCC in both *N*(*t*) and *ϕ*(*t*) greater than 0.8 (Supp Fig. S4B). From that filtered parameter set, we then found the Pareto boundary by removing any parameter sets where there existed another parameter set with a lower error in *N*(*t*) and *ϕ*(*t*) (Supp Fig. S4C). The resulting parameter sets formed a front, whereas we increase the regularization term *λ*, the error in *ϕ*(*t*) fit to the data decreased as the error in the *N*(*t*) data increased. With this set of Pareto front parameter sets, we looked at the individual parameter values and examined how they varied as we varied *λ* and thus as we improved the accuracy in fit to one data set over another (Supp Fig. S5). We could then look at the distribution of parameter values to see which parameter values were stable across objective function weightings, and which were most dependent on the weight of the data sets relative to one another (Fig 6L-Q, Supp Fig. S6). The parameter values chosen all fell well within the 95% CI of *θ*\* (Supp Fig S6).

#### Structural Identifiability of Model Parameters

We will demonstrate the structural identifiability of the individual model parameters using the differential algebra approach. Structural identifiability of a model and its parameters from a set of measurable outputs tells us that in theory, given perfect data, it is possible to uniquely identify model parameters. Structural identifiability is a pre-requisite for practical identifiability of model parameters from observed data. We start by presenting the non-dimensionalized model and measurement equations, assuming we can measure both N(t) and ϕ(t).

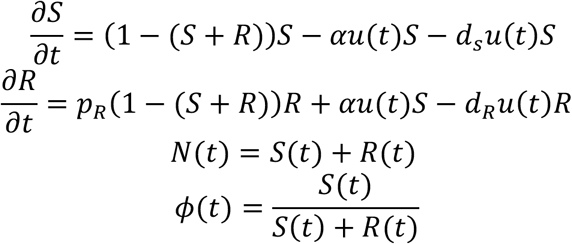

We assume all parameters are non-negative and 0 < *p_r_* < 1 represents the relative growth rate of the resistant population with respect to the sensitive population scaled by the carrying capacity, and *p_r_* < 1 assumes that resistant cells grow more slowly than sensitive cells. In work by Greene et al (16), they demonstrate that, if they assume d_r_ = 0, i.e. resistant cells are not killed by drug, and that the initial state of the population is completely comprised of sensitive cells (i.e. *N_0_*=*S_0_*), than the remaining parameters are uniquely identifiable from observations of total cell number alone.

We would like to extend this analysis by determining the identifiability of a new experimental system in which not only can *N*(*t*) = *S*(*t*) + *R*(*t*) be observed, but so also can the fraction of cells in each state over time, here denoted as *ϕ*(*t*). Under these circumstances, we want to test the identifiability of the model which now allows for a net-positive death rate due to drug, *d_R_*, and can have any composition of initial sensitive and resistant cells.

We follow the same arguments outlined in (16), along with the complete explanation of the approach with illustrative examples, for the case of multiple outputs from (62). We start by formulating the dynamical system relevant to our in vitro experimental system. Of note, even though we separately measure *N*(*t*) and *ϕ*(*t*) at discrete time points, since this analysis is for structural identifiability and assumes perfect, noise-free data, we will transform the observable outputs of *N*(*t*) and *ϕ*(*t*) into:

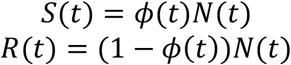

Treatment is initiated at time *t*=0, at which we make no assumptions about the composition of the population such that *S*(*0*) = *S_0_, R*(*0*) = *R_0_*. Here *0<S_0_+R_0_<1*. We note this is due to the non-dimensionalization in which we now track the proportion of confluent cells i.e. 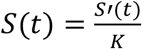 and 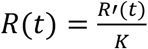 (see (16)) for additional details. We can now formulate our system in input/output form as:

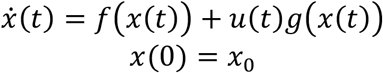

Where *f* and *g* are:

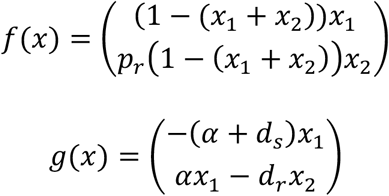

and *x*(*t*) = (*S*(*t*), *R*(*t*)). As is standard in control theory, the output is denoted by the variable y which in this work corresponds to *S*(*t*) and *R*(*t*) obtained from the transformations of the measured variables *N*(*t*) and *ϕ*(*t*)

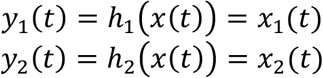

A system in this form is said to be uniquely structurally identifiable if the map (*p, u*(*t*)) → (*x*(*t, p*), *u*(*t*)) is injective (30,31,62), where p is the vector of parameters to be identified. In this instance *p* = (*S_0_, R_0_, d_s_, d_r_, α, p_r_*), the initial states and the parameters. Local identifiability and non-identifiability correspond to the map being finite-to-one and infinite-to-one, respectively. Our objective is then to demonstrate unique structural identifiability for model system and hence recover all parameter values p from the assumption of perfect, noise-free data.

To analyze identifiability, we utilize results appearing in (16) and (62), where a differential-geometric perspective is used. For the structural identifiability, we hypothesize that we have perfect (hence noise-free) input-output data is available of the form of *y_1_* and *y_2_* and its derivatives on any interval of time. We then, for example, make measurements of:

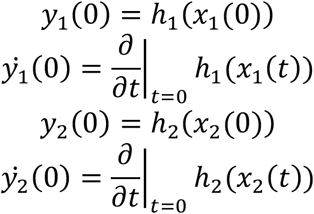

We can relate their values to the unknown parameter values p. If there exists inputs *u*(*t*) such that the above system of equations may be solved for p, the system is identifiable. The right-hand sides of the above the equation for *x*(*t*) may be computed in terms of the Lie derivatives of the vector fields *f* and *g*. The Lie differentiation *L_x_H* of a function *H* by a vector field *X* is given by:

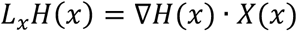

Iterated Lie derivatives are well-defined, and should be interpreted as the function composition, so that for example *L_y_L_x_H*(*x*) = L_y_(*L_x_H*) and 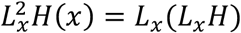.

Defining observable quantities at the zero-time derivatives of the generalized output *y* = *h*(*x*):

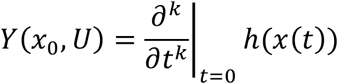

Where *U* ∈ *R^k^* is the value of the control *u*(*t*) and its derivatives evaluated at *t* = 0: *U* = (*u*(0), *u*′(0),… *u*^*k*−1^(0)). The initial conditions *x_0_* appear due to evaluation at *t*=0. The observation space is then defined as the span of the *Y*(*x*_0_, *U*) elements:

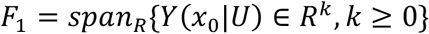

We also defined the span of iterated Lie derivatives with respect to the output vector fields *f*(*x*) and *g*(*x*):

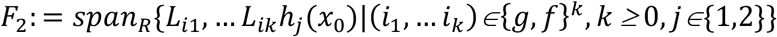

As is outlined in (62), (63) proved that *F_1_=F_2_*, so that the iterated Lie derivatives *F_2_* may be considered as the set of “elementary observables”. Hence, identifiability may be formulated in terms of the reconstruction of parameters *p* from elements in *F_2_*. Parameters p are then identifiable if the map

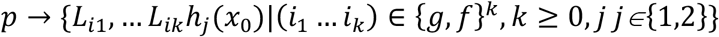

Is one-to-one. For the remainder of this analysis, we investigate the mapping defined here, because if one can reconstruct the values of *p* from the elementary observables (evaluated at the initial state), we can uniquely identify the parameters. This enables us to find the Lie derivatives for the two outputs *h_1_*(*x*) and *h_2_*(*x*), which will be found in terms of the parameters *p* and *x_1_* and *x_2_*. Then we can recall the evaluation at *t*=0 given by *x_0_* = (*S_0_, R_0_*), and our ability to observe these at *t*=0 allows us to set *x_1_* = *S_0_* and *x_2_* = *R_0_* and isolate the parameter *p* recursively from the observables and the Lie derivatives.

Using the input-output system written in terms of *f* and *g* we can write the following Lie derivatives:

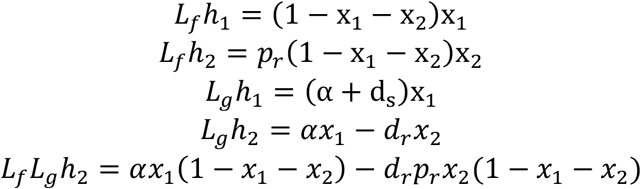

Recursively solving using *x_0_* = (*S_0_, R_0_*) to find the parameters *p*:

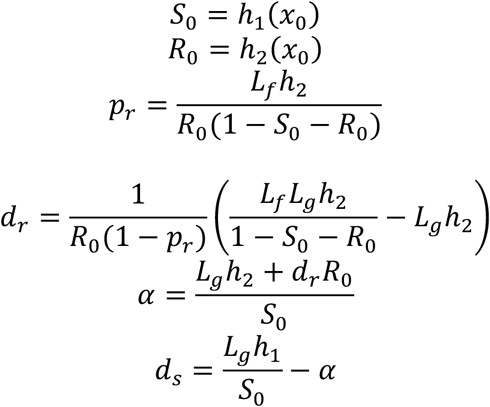

Since *F_1_* = *F_2_*, all of the above Lie derivatives are observable via appropriate treatment protocols. Thus by incorporating knowledge of *ϕ*(*t*), all parameters in system 1 are structurally identifiable. This represents an improvement over the identifiability with *N*(*t*) alone as a measurable output and allows us to introduce a non-zero *d_R_* parameter, which we have reason to believe based on experimental evidence, is the more biologically relevant scenario.

## Author Contributions

KJ and AB designed the study; GH, DM, EB, AG, and AA performed experiments; WM curated the data; KJ, GH, DM, EB, AG, RD and WM analyzed the data; KJ performed mathematical modeling; ES, AJ, TY advised on mathematical modeling, KJ and AB wrote the manuscript with input from all authors; all authors read and approved the manuscript.

## Acknowledgements

The authors are grateful for grant support from the NIH iMAT program (R21CA212928 to AB), CPRIT (RR1600005 to TEY), NCI (U01CA174706 and R01CA186193 to TEY) and NSF Grants #1716623 and #1849588 to EDS). KJ was supported by an NSF Graduate Research Fellowship 1610403. T.E.Y. is a CPRIT Scholar of Cancer Research. The authors also thank the Genomic and Sequencing Analysis Facility at the University of Texas at Austin and Dennis Wylie for advice throughout the project.

## References

1. Ferrall-Fairbanks MC, Ball M, Padron E, Altrock PM. Leveraging Single-Cell RNA Sequencing Experiments to Model Intratumor Heterogeneity. Clin Cancer Informatics [Internet]. 2019; 1–10. Available from: https://doi.org/10.1200/CCI.18.00074

2. Syed AK, Woodall R, Whisenant JG, Yankeelov TE, Sorace AG. Characterizing Trastuzumab-Induced Alterations in Intratumoral Heterogeneity with Quantitative Imaging and Immunohistochemistry in HER2 + Breast Cancer. Neoplasia [Internet]. 2019;21(1):17–29. Available from: https://doi.org/10.1016/j.neo.2018.10.008

3. Pyne S, Hu X, Wang K, Rossin E, Lin T, Maier LM, et al. Automated highdimensional flow cytometric data analysis. Proc Natl Acad Sci [Internet]. 2009;106(21):8519–24. Available from: www.pnas.org/cgi/doi/10.1073/pnas.0903028106

4. Islam S, Zeisel A, Joost S, La Manno G, Zajac P, Kasper M, et al. Quantitative single-cell RNA-seq with unique molecular identifiers. Nat Methods. 2014;11(2).

5. Guo J, Grow EJ, Yi C, Micochova H, Maher GJ, Lindskog C, et al. Chromatin and Single-Cell RNA-Seq Profiling Reveal Dynamic Signaling and Metabolic Transitions during Human Spermatogonial Stem Cell Development. Cell Stem Cell [Internet]. 2018;21(4):533–46. Available from: https://doi.org/10.1016/j.stem.2017.09.003

6. Kumar MP, Du J, Lagoudas G, Yang J, Sawyer A, Drummond DC, et al. Analysis of Single-Cell RNA-Seq Identifies Cell-Cell Communication Associated with Tumor Characteristics. Cell Rep [Internet]. 2018;25(6):1458-1468.e4. Available from: https://doi.org/10.1016/j.celrep.2018.10.047

7. Wang Y, Wang R, Zhang S, Song S, Jiang C, Han G, et al. iTALK: an R Package to Characterize and Illustrate Intercellular Communication. bioRxiv [Internet]. 2019; Available from: https://dx.doi.org/10.1101/507871

8. Zhao X, Wu S, Fang N, Sun X, Fan J. Evaluation of single-cell classifiers for single-cell RNA sequencing data sets. 2019;00(July):1–15.

9. Cho H, Ayers K, Depills L, Park J, Radunskaya A, Rockne R. Modelling acute myeloid leukaemia in a continuum of differentiation states. Lett Biomath. 2018;5:1–41.

10. Manno G La, Soldatov R, Zeisel A, Braun E, Hochgerner H, Petukhov V, et al. RNA velocity of single cells. Nature [Internet]. 2018; Available from: http://dx.doi.org/10.1038/s41586-018-0414-6

11. Al’Khafaji A, Gutierrez C, Brenner E, Durrett R, Johnson KE, Zhang W, et al. Expressed barcodes enable clonal characterization of chemotherapeutic responses in chronic lymphocytic leukemia. bioRxiv [Internet]. 2019;1–24. Available from: https://dx.doi.org/10.1101/761981

12. Smalley I, Kim E, Li J, Spence P, Wyatt CJ, Eroglu Z, et al. Leveraging transcriptional dynamics to improve BRAF inhibitor responses in melanoma. EBioMedicine [Internet]. 2019; Available from: https://doi.org/10.1016/j.ebiom.2019.09.023

13. Stumpf PS, Smith RCG, Lenz M, Schuppert A, Müller FJ, Babtie A, et al. Stem Cell Differentiation as a Non-Markov Stochastic Process. Cell Syst. 2017;5(3):268–282.e7.

14. Brady R, Nagy JD, Gerke TA, Zhang T, Wang AZ, Zhang J, et al. Prostate-Specific Antigen Dynamics Predict Individual Responses to Intermittent Androgen Deprivation. bioRxiv [Internet]. 2019; Available from: https://dx.doi.org/10.1101/624866

15. McKenna MT, Weis JA, Quaranta V, Yankeelov TE. Variable Cell Line Pharmacokinetics Contribute to Non-Linear Treatment Response in Heterogeneous Cell Populations. Ann Biomed Eng [Internet]. 2018/02/26. 2018 Jun;46(6):899–911. Available from: https://www.ncbi.nlm.nih.gov/pubmed/29484528

16. Greene JM, Sanchez-Tapia C, Sontag ED. Mathematical Details on a Cancer Resistance Model. bioRxiv [Internet]. 2018;1–42. Available from: https://dx.doi.org/10.1101/475533

17. Greene JM, Gevertz JL, Sontag ED. Mathematical Approach to Differentiate Spontaneous and Induced Evolution to Drug Resistance During Cancer Treatment abstract. JCO Clin Cancer Informatics. 2019;42–9.

18. Gevertz JL, Greene JM, Sontag ED. Validation of a Mathematical Model of Cancer Incorporating Spontaneous and Induced Evolution to Drug Resistance. bioRxiv [Internet]. 2019;1–15. Available from: http://dx.doi.org/10.1101/2019.12.27.889444

19. Gatenby RA, Silva AS, Gillies RJ, Frieden BR. Adaptive therapy. Cancer Res. 2009;69(11):4894–903.

20. Prokopiou S, Moros EG, Poleszczuk J, Caudell J, Torres-roca JF, Latifi K, et al. A proliferation saturation index to predict radiation response and personalize radiotherapy fractionation. Radiat Oncol [Internet]. 2015;1–8. Available from: http://dx.doi.org/10.1186/s13014-015-0465-x

21. Howard GR, Johnson KE, Ayala AR, Yankeelov TE, Brock A. A multi-state model of chemoresistance to characterize phenotypic dynamics in breast cancer. Sci Rep [Internet]. 2018;(July):1–11. Available from: http://dx.doi.org/10.1038/s41598-018-30467-w

22. McKenna MT, Weis JA, Quaranta V, Yankeelov TE. Variable Cell Line Pharmacokinetics Contribute to Non-Linear Treatment Response in Heterogeneous Cell Populations. Ann Biomed Eng. 2018;46(6):899–911.

23. McKenna MT, Weis JA, Barnes SL, Tyson DR, Miga MI, Quaranta V, et al. A Predictive Mathematical Modeling Approach for the Study of Doxorubicin Treatment in Triple Negative Breast Cancer. Sci Rep [Internet]. 2017;7(1):1–14. Available from: http://dx.doi.org/10.1038/s41598-017-05902-z

24. Johnson KE, Gomez A, Burton J, White D, Chakravarty A, Schmid A, et al. Directional inconsistency between Response Evaluation Criteria in Solid Tumors (RECIST) time to progression and response speed and depth. Eur J Cancer [Internet]. 2019;109:196–203. Available from: https://doi.org/10.1016/j.ejca.2018.11.008

25. Al’Khafaji AM, Deatherage D, Brock A. Control of Lineage-Specific Gene Expression by Functionalized gRNA Barcodes. ACS Synth Biol. 2018;

26. Shin VY, Chen J, Cheuk IWY, Siu MT, Ho CW, Wang X, et al. Long non-coding RNA NEAT1 confers oncogenic role in triple-negative breast cancer through modulating chemoresistance and cancer stemness. Cell Death Dis [Internet]. 2019;10(4). Available from: http://dx.doi.org/10.1038/s41419-019-1513-5

27. Censor Y. Pareto Optimality in Multiobjective Problems. Appl Math Optim. 1977;4:41–59.

28. Thermo Fisher Scientific. Useful Numbers for Cell Culture [Internet]. [cited 2020 Feb 11]. Available from: https://www.thermofisher.com/us/en/home/references/gibco-cell-culture-basics/cell-culture-protocols/cell-culture-useful-numbers.html

29. Eisenberg MC, Robertson SL, Tien JH. Identifiability and estimation of multiple transmission pathways in cholera and waterborne disease. J Theor Biol [Internet]. 2013;324:84–102. Available from: http://dx.doi.org/10.1016/j.jtbi.2012.12.021

30. Eisenberg MC. Input-output equivalence and identifiability: some simple generalizations of the differential algebra approach. arXiv. 2019;1–25.

31. Brouwer AF, Meza R, Eisenberg MC, Arbor A. A systematic approach to determining the identifiability of multistage carcinogenesis models. Risk Anal. 2017;37(7):1375–87.

32. Raue A, Kreutz C, Maiwald T, Bachmann J, Schilling M, Klingmüller U, et al. Structural and practical identifiability analysis of partially observed dynamical models by exploiting the profile likelihood. Bioinformatics. 2009;25(15):1923–9.

33. Suvà ML, Tirosh I. Single-Cell RNA Sequencing in Cancer: Lessons Learned and Emerging Challenges. Mol Cell. 2019;75(1):7–12.

34. Levitin HM, Yuan J, Sims PA. Single-Cell Transcriptomic Analysis of Tumor Heterogeneity. Trends in Cancer. 2018 Apr;4(4):264–8.

35. Rockne RC, Hawkins-daarud A, Swanson KR, Sluka JP, Glazier JA, Macklin P, et al. The 2019 mathematical oncology roadmap The 2019 mathematical oncology roadmap. 2019;

36. McKenna MT, Weis JA, Brock A, Quaranta V, Yankeelov TE. Precision Medicine with Imprecise Therapy: Computational Modeling for Chemotherapy in Breast Cancer. Transl Oncol [Internet]. 2018;11(3):732–42. Available from: https://doi.org/10.1016/j.tranon.2018.03.009

37. Jarrett AM, Lima EABF, Hormuth DA, McKenna MT, Feng X, Ekrut DA, et al. Mathematical models of tumor cell proliferation: A review of the literature. Expert Rev Anticancer Ther [Internet]. 2018 Dec 2;18(12):1271–86. Available from: https://doi.org/10.1080/14737140.2018.1527689

38. Poleszczuk J, Enderling H. The Optimal Radiation Dose to Induce Robust Systemic Anti-Tumor Immunity. Int J Mol Sci. 2018; 19(11).

39. Zhang Y, Huynh JM, Liu G, Ballweg R, Aryeh KS, Paek AL, et al. Designing combination therapies with modeling chaperoned machine learning. PLoS Comput Biol [Internet]. 2019; 15(9):1–17. Available from: http://dx.doi.org/10.1371/journal.pcbi.1007158

40. Badri H, Pitter K, Holland EC, Michor F, Leder K. Optimization of radiation dosing schedules for proneural glioblastoma. J Math Biol. 2016;72(5):1301–36.

41. Poleszczuk J, Enderling H, Poleszczuk J, Enderling H. Cancer Stem Cell Plasticity as Tumor Growth Promoter and Catalyst of Population Collapse. Stem Cells Int [Internet]. 2016;2016:1–12. Available from: http://www.hindawi.com/journals/sci/2016/3923527/

42. Greene JM, Levy D, Fung KL, Souza PS, Gottesman MM, Lavi O. Modeling intrinsic heterogeneity and growth of cancer cells. J Theor Biol [Internet]. 2015;367:262–77. Available from: http://dx.doi.org/10.1016/j.jtbi.2014.11.017

43. Jarrett AM, Bloom MJ, Ekrut DA, Yankeelov TE. Mathematical modelling of trastuzumab-induced immune response in an in vivo murine model of HER2 + breast cancer. 2018;2:1–30.

44. Hormuth DA, Jarrett AM, Feng X, Yankeelov TE. Calibrating a Predictive Model of Tumor Growth and Angiogenesis with Quantitative MRI. Ann Biomed Eng. 2019;47(7):1539–51.

45. Yankeelov TE, Atuegwu N, Hormuth D, Weis JA, Barnes SL, Miga MI, et al. Clinically Relevant Modeling of Tumor Growth and Treatment Response. 2013;5(187):1–6.

46. Yankeelov TE, Quaranta V, Evans KJ, Rericha EC. Toward a science of tumor forecasting for clinical oncology. Cancer Res. 2015;75(6):918–23.

47. Ma K-Y, Schonnesen AA, Brock A, Van Den Berg C, Eckhardt SG, Liu Z, et al. Single-cell RNA sequencing of lung adenocarcinoma reveals heterogeneity of immune response–related genes. JCI Insight [Internet]. 2019 Feb 21;4(4). Available from: https://doi.org/10.1172/jci.insight.121387

48. Luecken MD, Theis FJ. Current best practices in single-cell RNA-seq analysis: a tutorial. Mol Syst Biol. 2019;15(e8746).

49. He J, Ahuja N. Personalized Approaches to Gastrointestinal Cancers: Importance of Integrating Genomic Information to Guide Therapy. Surg Clin North Am. 2015;95(5):1081–94.

50. Kowarz E, Loescher D, Marschalek R. Optimized Sleeping Beauty transposons enable robust stable transgenic cell lines. Biotechnol J. 2015;41:647–53.

51. Mátés L, Chuah MKL, Belay E, Jerchow B, Manoj N, Acosta-Sanchez A, et al. Molecular evolution of a novel hyperactive Sleeping Beauty transposase enables robust stable gene transfer in vertebrates. Nat Genet. 2009;41(6):753–61.

52. Wolf FA, Angerer P, Theis FJ. SCANPY: large-scale single-cell gene expression data analysis. Genome Biol [Internet]. 2018; 19(1):15. Available from: https://doi.org/10.1186/s13059-017-1382-0

53. Büttner M, Miao Z, Wolf FA, Teichmann SA, Theis FJ. A test metric for assessing single-cell RNA-seq batch correction. Nat Methods [Internet]. 2019;16(1):43–9. Available from: https://doi.org/10.1038/s41592-018-0254-1

54. Vieth B, Parekh S, Ziegenhain C, Enard W, Hellmann I. A systematic evaluation of single cell RNA-seq analysis pipelines. Nat Commun [Internet]. 2019;10(1):4667. Available from: https://doi.org/10.1038/s41467-019-12266-7

55. Traag VA, Waltman L, van Eck NJ. From Louvain to Leiden: guaranteeing well-connected communities. Sci Rep [Internet]. 2019;9(1):5233. Available from: https://doi.org/10.1038/s41598-019-41695-z

56. L. Lun AT, Bach K, Marioni JC. Pooling across cells to normalize single-cell RNA sequencing data with many zero counts. Genome Biol [Internet]. 2016;17(1):75. Available from: https://doi.org/10.1186/s13059-016-0947-7

57. Tirosh I, Izar B, Prakadan SM, Ii MHW, Treacy D, Trombetta JJ, et al. Dissecting the multicellular ecosystem of metastatic melanoma by single-cell RNA-seq. Science (80-). 2019;352(6282).

58. Tunio IA, Soomro S, Soomro TA, Bhatti MT, Shaikh M. Face Recognition System by using Eigen Value Decomposition. Int J Comput Sci Netw Secur. 2018;18(5):8–12.

59. Anderson ARA, Weaver AM, Cummings PT, Quaranta V. Tumor Morphology and Phenotypic Evolution Driven by Selective Pressure from the Microenvironment. Cell. 2006; 127(5):905–15.

60. Jarrett AM, Liu Y, Cogan NG, Hussaini MY. Global sensitivity analysis used to interpret biological experimental results. J Math Biol [Internet]. 2015;71:151–70. Available from: http://dx.doi.org/10.1007/s00285-014-0818-3

61. Coetzee FM, Stonick VL. On a natural homotopy between linear and nonlinear single-layer networks. IEEE Trans Neural Networks. 1996;7(2):307–17.

62. Sontag ED. Dynamic compensation, parameter identifiability, and equivariances. PLoS Comput Biol. 2017;13(4):1–17.

63. Wang Y, Sontag ED. On two definitions of observation spaces. Syst Control Lett. 1989;13:213–8.

