## Supplementary figures for "Integrating multimodal data sets into a mathematical framework to describe and predict therapeutic resistance in cancer"

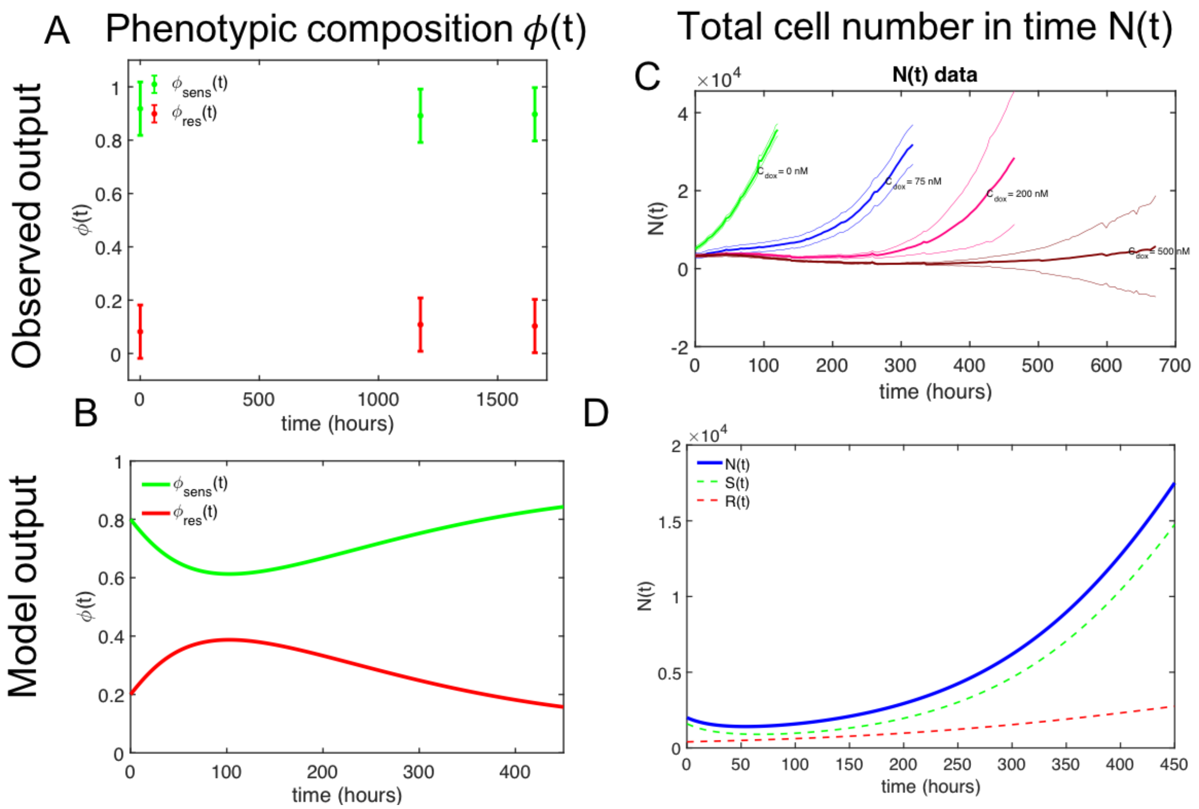


**Supplementary Figure S1. Measured and model predicted outputs to be used for parameter estimation from observed data** A. Observed estimated fraction of sensitive cells (green) and resistant cells (red) from scRNAseq classifier at three time points $\phi$(t). B. Model predicted output of sensitive cell fraction dynamics (green) and resistant cell fraction dynamics (red) for an example parameter set. C. Observed number of tumor cells in time for pulse treatments of doxorubicin at 0, 75, 200, and 500 nM. D. Model predicted output of total cell number in time for a pulse treatment of 75 nM for an example parameter set.

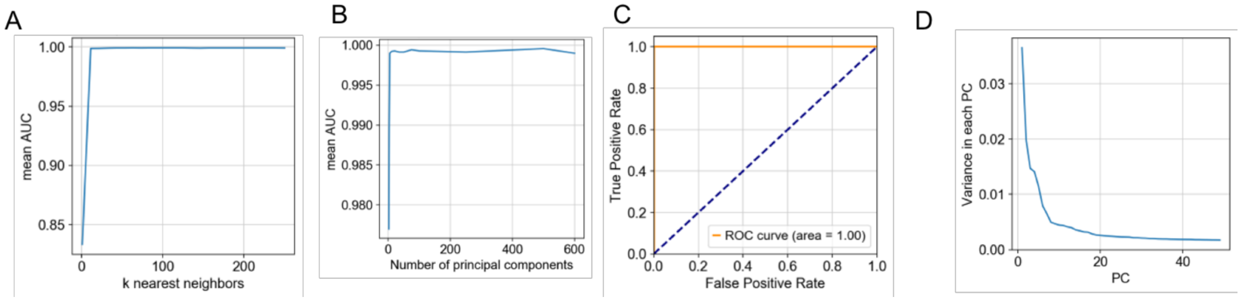


**Supplementary Figure S2. Optimization of Principal Component Classifier Hyperparameters use coordinate optimization and 5-fold Cross Validation.** A. Number of nearest neighbors used in the classifier versus mean AUC from 5-fold CV to determine optimal number of neighbors of k=73. B. Number of principal components used in the classifier versus mean AUC from 5-fold CV to determine optimal number of components, n=500. C. ROC curve from classifier with optimized number of nearest neighbors and components for separating labeled cells. D. Proportion of variance explained by the principal components drops off sharply for higher PCs.

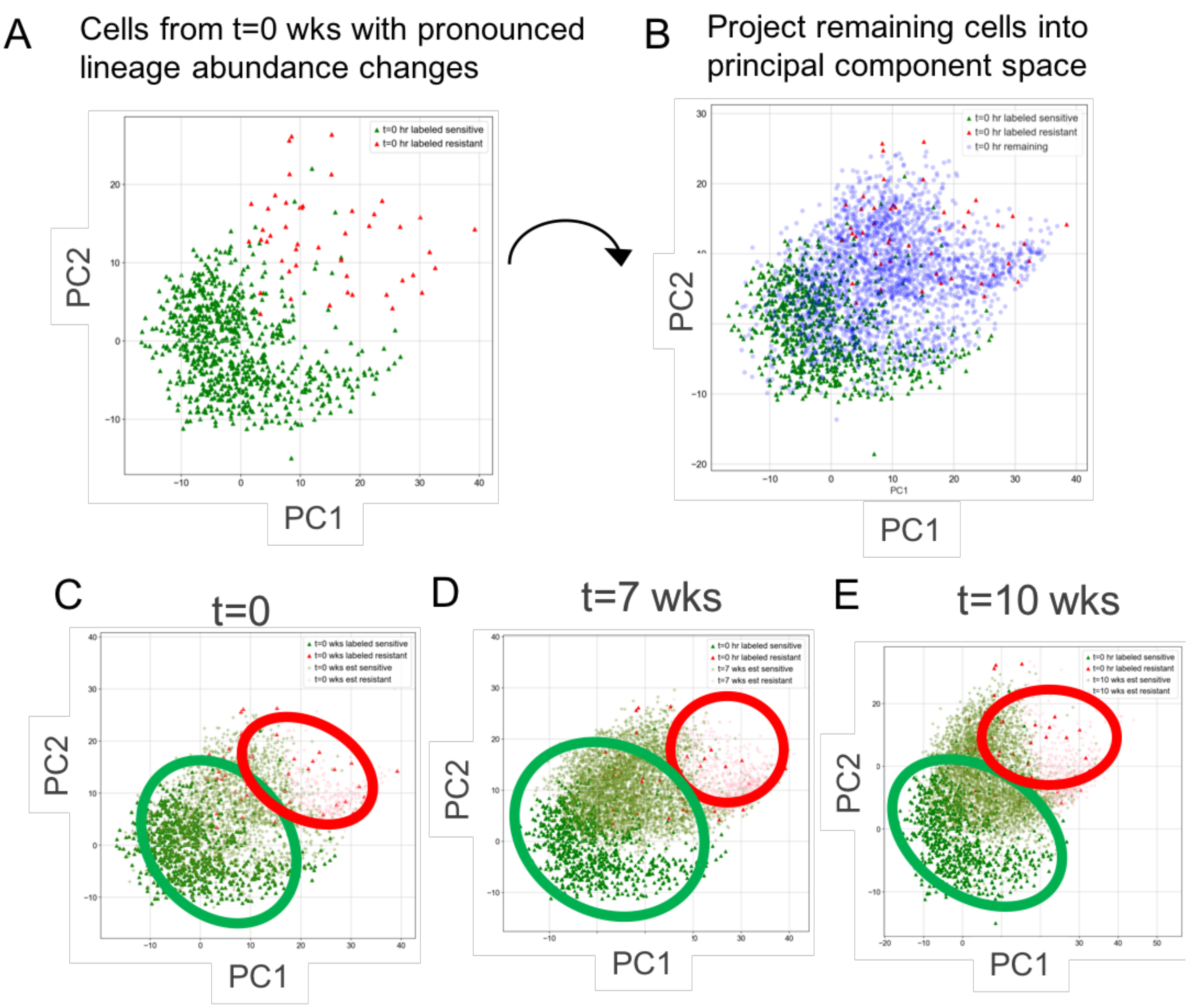


**Supplementary Figure S3. Single cell transcriptomes from each time point projected into principal component space and classified using nearest neighbors** A. Lineage-abundance guided "labeled" cells projected into principal component space separate along components (PC1 and PC2 shown here for visual effect). B. Unknown cells are projected into the principal component space of the labeled cells. C. Remaining cells from t=0 projected onto labeled cells in PC space and estimated as sensitive (olive) or resistant (green). D. Cells from t=7 weeks projected alongside labeled cells. E. Cells from t=10 weeks projected alongside labeled cells.

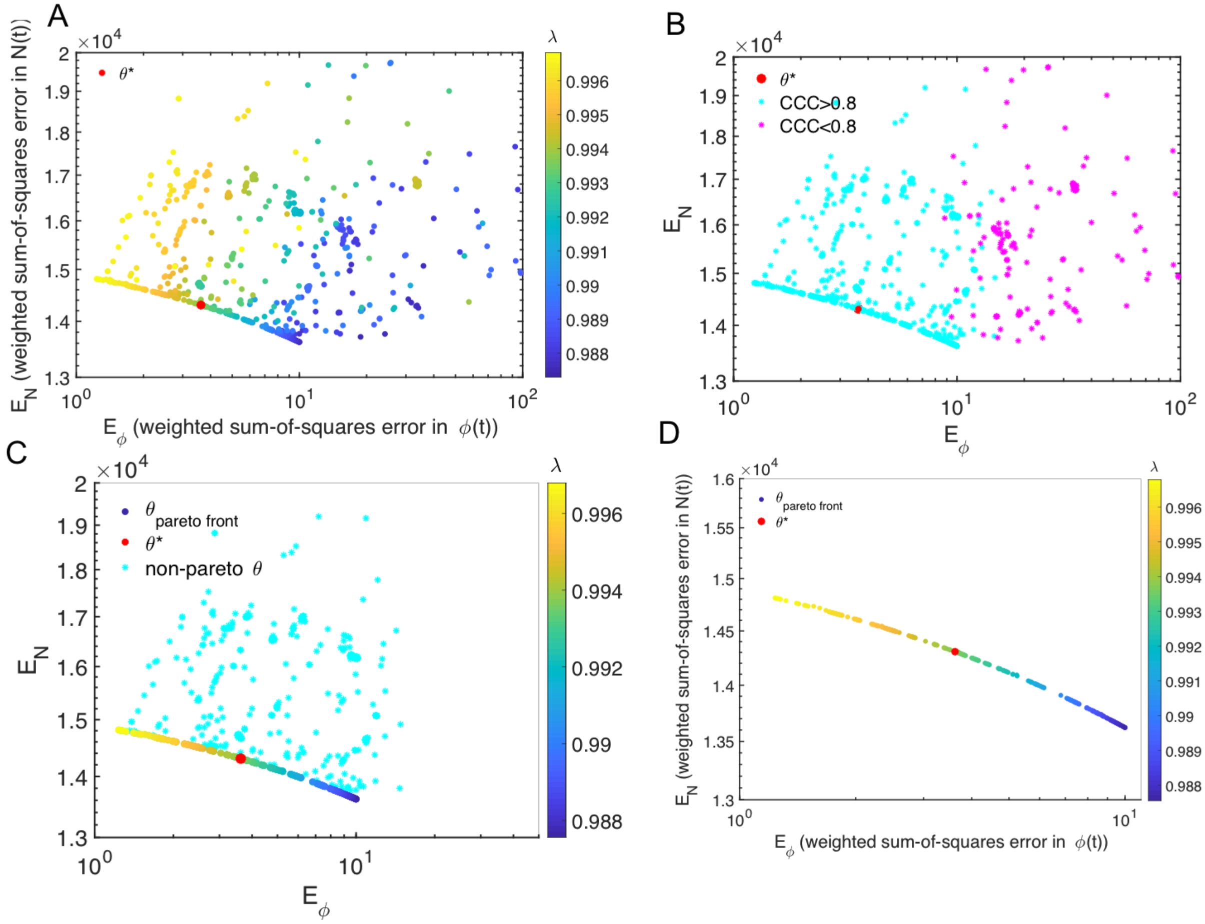


**Supplementary Figure S4. Step-by-step refinement of accepted parameter sets to identify solutions along pareto front.** A. The resulting weighted sum-squared error in N(t) ($E_N$) and $\phi$(t) ($E\phi$ ) of 1000 optimizations with the regularization term $\boldsymbol{\lambda}$, varying from $\boldsymbol{\lambda}$=0 (only fitting N(t) data), to $\boldsymbol{\lambda}$=1 (only fitting $\phi$(t) data). B. Filtering of parameter sets to require that parameter sets have a CCC>0.8 in both N(t) and $\phi$(t). C. Further filtering of parameter sets to remove "non-pareto" solutions- i.e. any parameter sets theta for which there exists another theta with a lower error in both $\phi$(t) and N(t). D. The final set of "pareto-front" solutions, which contain parameter sets for which an improvement in error in N(t) comes at a trade-off of a worsening in error in $\phi$(t).

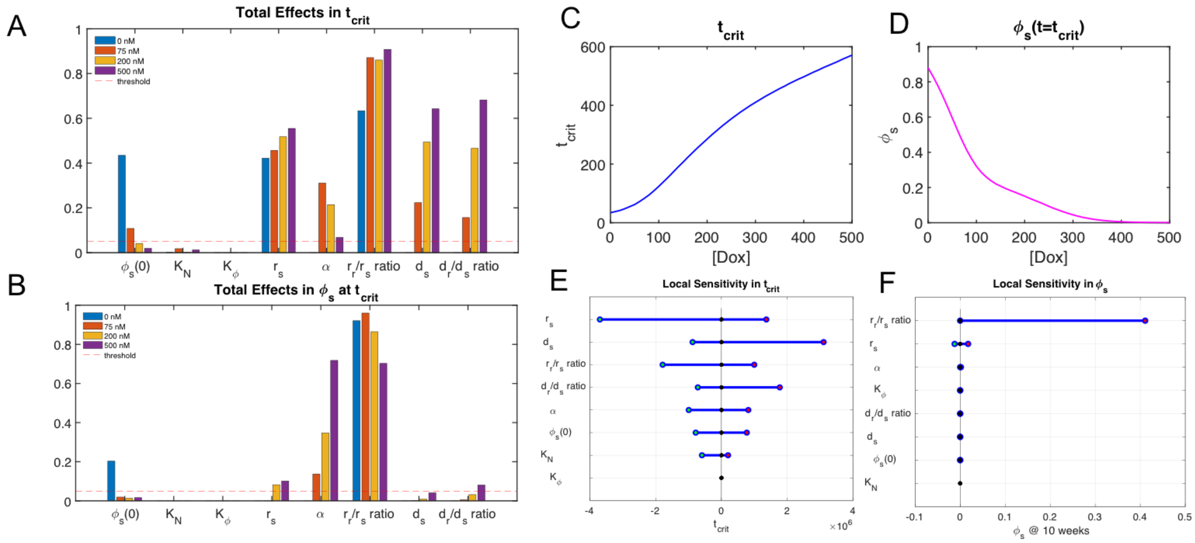


**Supplementary Figure S5. Sensitivity Analysis of Model Parameters Reveals All Parameters are Locally and Globally Sensitive Under Treatment.** A. Sobol's total effects of each parameter globally on critical time for 0,75, 200, and 500 nM pulse treatments reveals that all fit parameters are above the threshold of sensitivity for at least one of those doses (the parameter contributes at least 5% to the critical time for at least one of the doxorubicin concentrations). B. Sobol's total effects of each parameter globally on sensitive cell fraction for 0, 75, 200 and 500 nM pulse treatments reveals that most fit parameters are above the threshold of sensitivity for at least one of the doses. The carrying capacity of the single cell RNA sequencing experiment ($K_2$) is the only parameter that is not above the threshold for any sensitivity analysis output or dose, and for this reason supports our decision to set that carrying capacity from a literature value (the expected number of 231 cells at confluence in a 10 cm dish, which the cells were expanded up to). C. An example of the model predicted critical time as a function of doxorubicin concentration, taken from the selected parameter set in red in Fig 5A. Critical time is chosen as an output for model sensitivity because it evaluates treatment response and drug sensitivity in of a cell population:drug concentration combination without biasing for response dynamics that might vary from system to system, and because it is most relevant to what we experimentally are able to observe (i.e. the cells rebounded to 2 times their initial cell number on this day). D. An example of the model predicted sensitive cell fraction at the critical time as a function of doxorubicin concentration, again for the selected parameter set in red in Fig 5A. This was chosen again because of its relevance to experimental workflows, as the time at which the population rebounds to 2 the seeding population might be a good time at which we could perform an experimental analysis of the tumor cell composition (i.e. scRNAseq). E. Local sensitivity in critical time produced by varying the selected parameter set by 50% above and below its value and recording the resulting change in critical time trajectory over a doxorubicin range of 0 to 500 nM. F. Local sensitivity in sensitive cell fraction at critical time produced by again varying the selected parameter set by 50% above and below its value and recording the resulting change in sensitive cell fraction over a doxorubicin range of 0 to 500 nM.

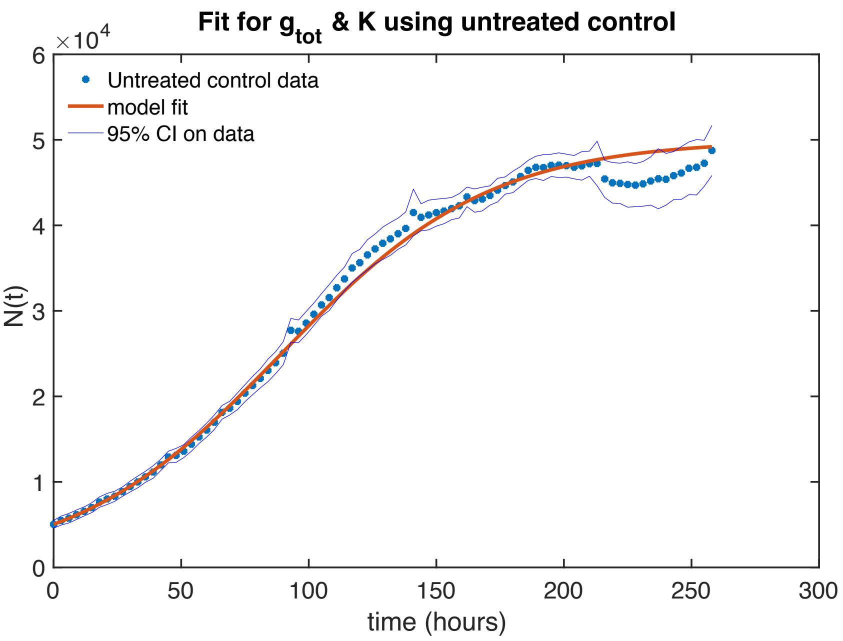


**Supp. Fig. S6. Fit to untreated control** to find effective dose and carrying capacity of MDA-MB- 231 cells in a 96 well plate.

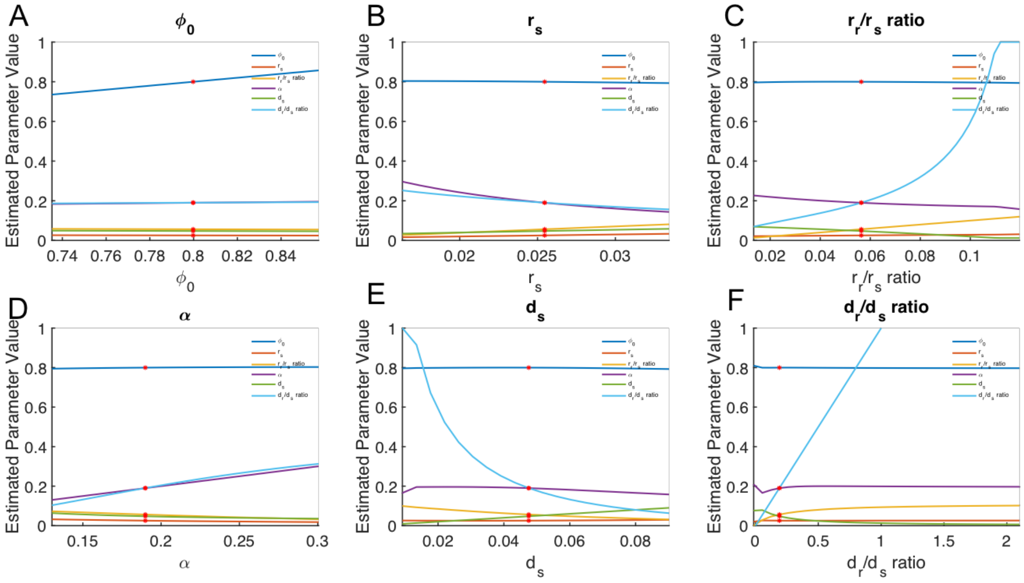


**Supp Fig S7. Parameter relationships from profile likelihood analysis.** Plot of the how each of the remaining 5 parameters varied while “profiling” A. $\phi_0$, B. $r_s$ C. $r_r/r_s$ ratio D. α, E. $d_s$, and F. $d_r/d_s$ ratio. The resulting curves describe the parameter relationships that enable the model to fit the observed data set.

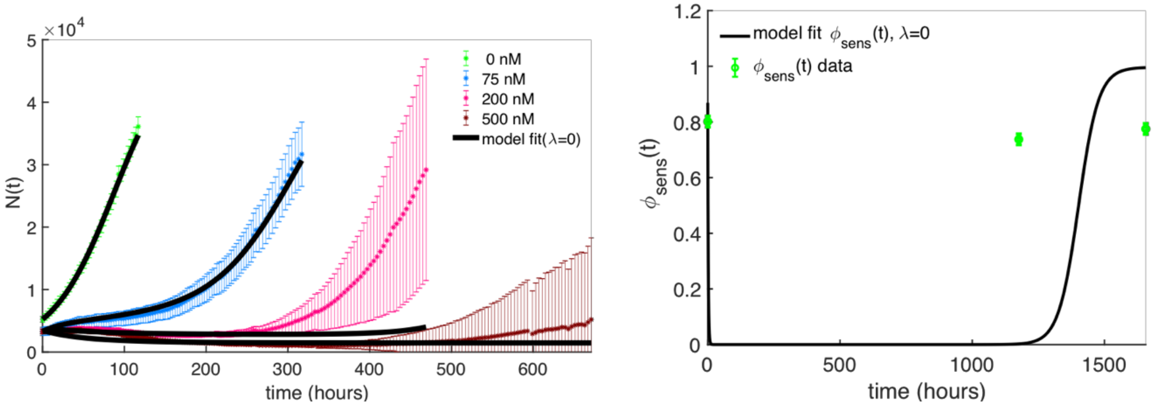


**Supplementary Figure S8. Fitting results without incorporating $\phi$(t).** A. Model fit compared to N(t) when only using N(t) data for calibration. Because the higher doses (200 nM and 500 nM) have a higher data uncertainty, we do not fit these doses well. The CCC of the mean N(t) compared to the model calibrated N(t) is 0.8471. B. Resulting model prediction of $\phi$(t) dynamics, based on calibration from N(t) data only, with a CCC of 0.0913.

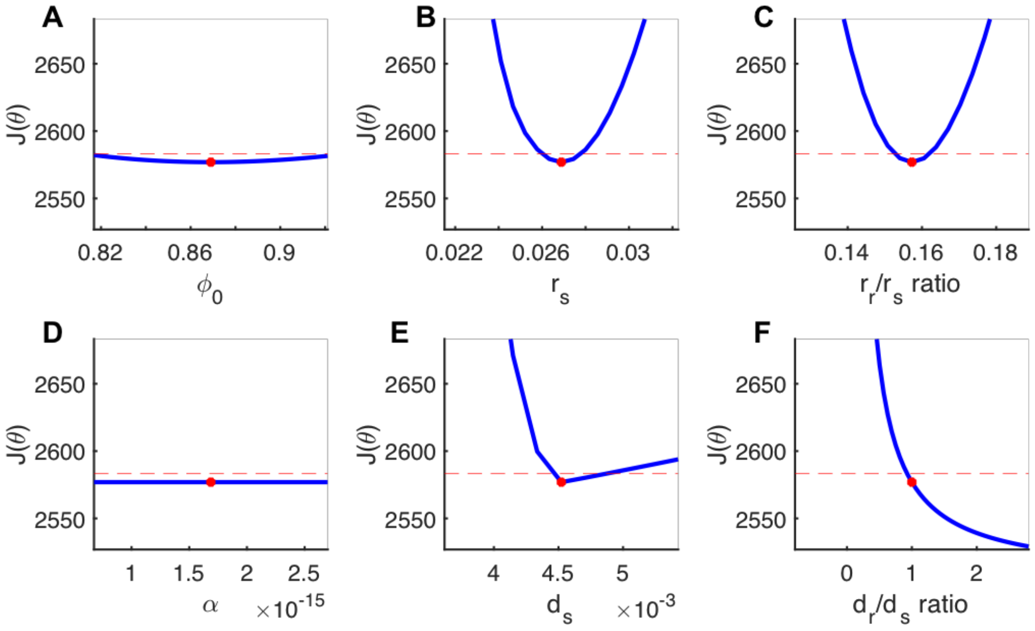


**Supplementary Figure S9. Profile likelihood results reveal that not all parameter are identifiable without incorporating $\phi$(t).** A. Profile likelihood around $\phi_0$, the initial sensitive cell fraction, reveals that in this case, parameter is identifiable, as it does eventually cross the 95% $\chi^2$- threshold. B. Profile likelihood around the sensitive cell growth rate, $r_s$, revealing the parameter is identifiable. C. Profile likelihood around the resistant-to-sensitive cell growth rate ratio reveals the parameter is identifiable. D. Profile likelihood around the drug-induced resistance rate α, revealing the parameter is unidentifiable because none of the profiled values enable the objective function value, J(θ) to cross above the threshold, indicating the value of this parameter within this region does not affect the goodness of fit of the model to the data. E. Profile likelihood around the sensitive cell death rate, $d_s$, revealing the parameter is identifiable. F. Profile likelihood around the resistant-to-sensitive cell death rate ratio, revealing the parameter is unidentifiable at the upper bound because the profiled values do not cross the objective function threshold, and therefore their value cannot be uniquely identified.

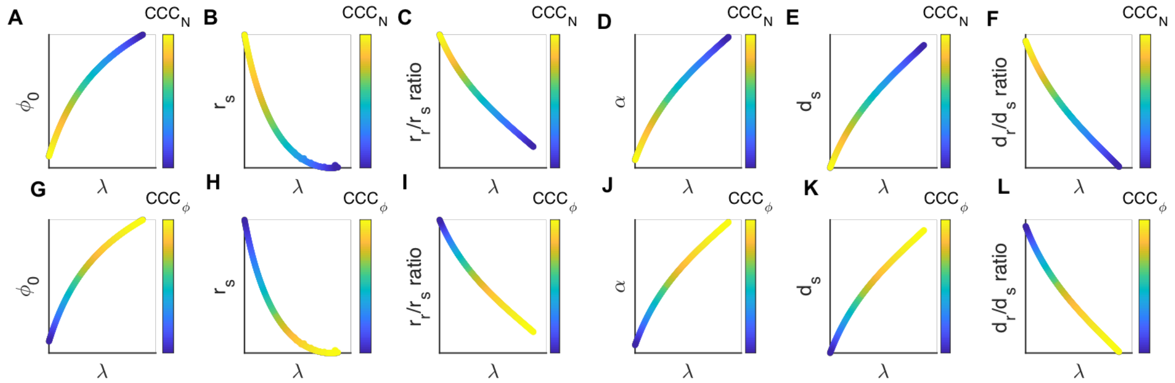


**Supplementary Figure S10. Variation of parameter values as a function of the regularization term λ, indicate that parameter values have directional bias in their goodness of fit in N(t) vs. the $\phi$(t).** A-F. Parameter values over the range of λ in the pareto front, colored by the value's corresponding accuracy in calibration to the N(t) data ($CCC_N$). G-L. Parameter values over the range of lambda in the pareto front, colored by the value's corresponding accuracy in calibration to the $\phi$(t) data ($CCC_\phi$).

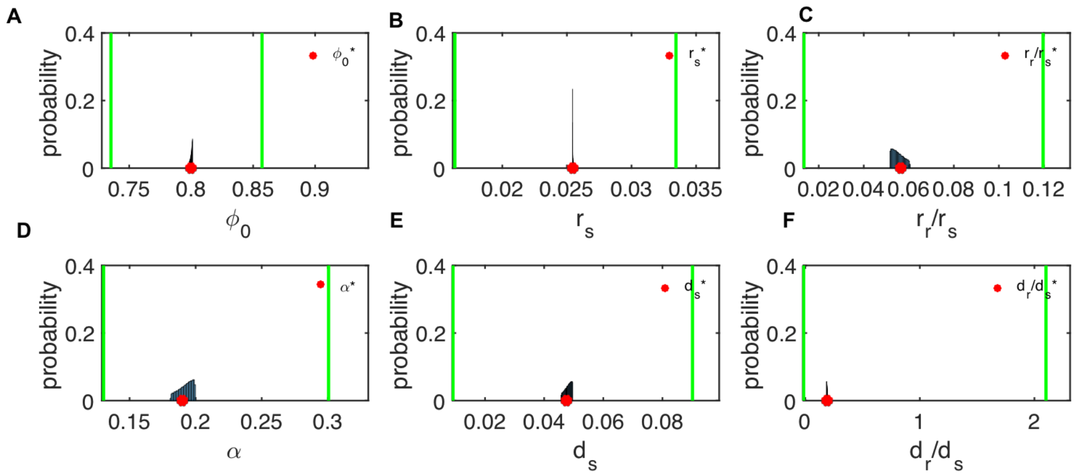


**Supplementary Figure S11. Pareto front parameter distributions fall well within the 95 % CI on θ*, found via the prolife likelihood method (Fig 6F-K) and displayed here by the green lines.** A. Distribution of pareto front accepted parameter $\phi_0$. B. Distribution of pareto front accepted parameter $r_s$. C. Distribution of pareto front accepted parameter resistant to sensitive growth rate. D. Distribution of pareto front accepted parameter α. E. Distribution of pareto front accepted parameter $d_s$. F. Distribution of pareto front accepted parameter resistant to sensitive cell death rate.

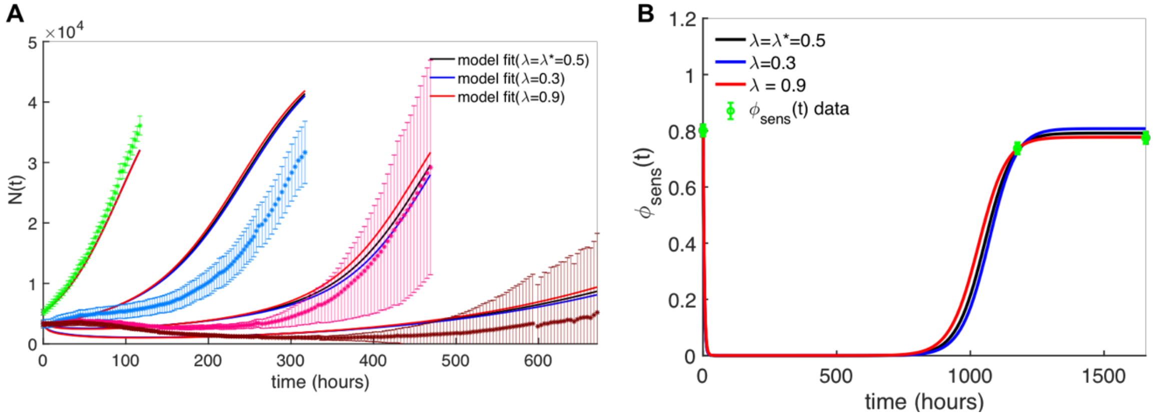


**Supp Fig S12. Fitting results for the range of pareto-front parameter sets** A. Model fit to N(t) for the weighting by number of data points only (black, **λ** = **λ***), the lowest **λ** (**λ** = 0.3) favoring N(t) the most, and the highest **λ** (**λ**= 0.9) favoring $\phi$(t) the most. B Model fit to $\phi$(t) for the range of accepted pareto front parameter sets.

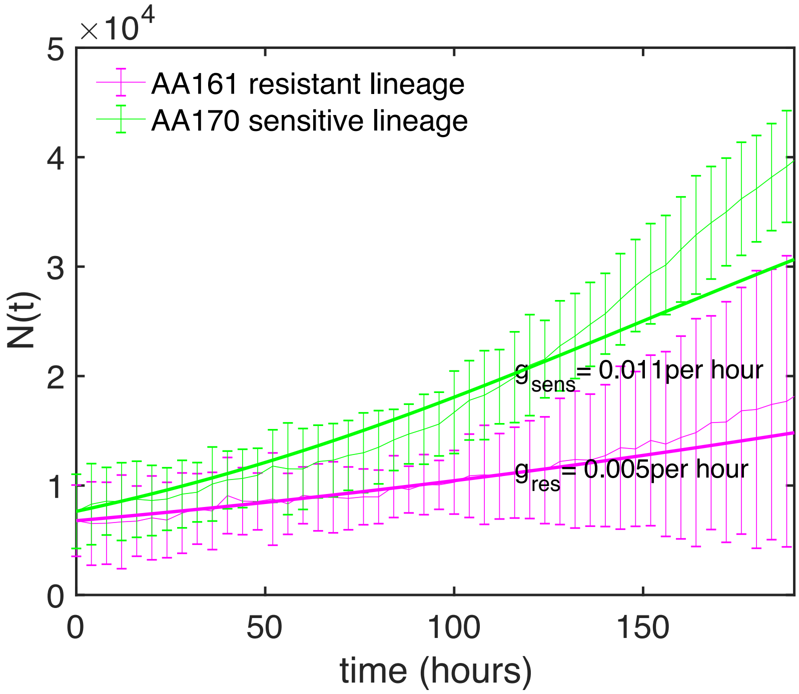


**Supplementary Figure S13. Growth dynamics of isolated sensitive and resistant cell lineages** indicates that sensitive cells growth on more quickly than the resistant cells, validating our modeling assumptions.
